# Bioreversible anionic cloaking enables intracellular protein delivery with ionizable lipid nanoparticles

**DOI:** 10.1101/2024.01.20.576479

**Authors:** Azmain Alamgir, Souvik Ghosal, Matthew P. DeLisa, Christopher A. Alabi

## Abstract

Protein-based therapeutics comprise a rapidly growing subset of pharmaceuticals, but enabling their delivery into cells for intracellular applications has been a longstanding challenge. To overcome the delivery barrier, we explored a reversible, bioconjugation-based approach to modify the surface charge of protein cargos with an anionic “cloak” to facilitate electrostatic complexation and delivery with lipid nanoparticle (LNP) formulations. We demonstrate that the conjugation of lysine-reactive sulfonated compounds can allow for the delivery of various protein cargos using FDA-approved LNP formulations of the ionizable cationic lipid DLin-MC3-DMA (MC3). We apply this strategy to functionally deliver RNase A for cancer cell killing as well as a full-length antibody to inhibit oncogenic β-catenin signaling. Further, we show that LNPs encapsulating cloaked fluorescent proteins distribute to major organs in mice following systemic administration. Overall, our results point towards a generalizable platform that can be employed for intracellular delivery of a wide range of protein cargos.

## Introduction

Proteins possess a remarkable capacity to execute a wide array of intricate functions in biology. Given their diverse roles, proteins have been extensively explored as potential therapeutic agents for addressing human diseases. In contrast to small-molecule drugs that have long dominated the pharmacopeia, protein-based therapeutics can offer reduced toxicity, enhanced bioavailability, and highly specific modes of biological activity.^1^ The growth of protein therapies in the clinic has been remarkable; in 2022 alone, nearly half of all FDA-approved drugs consisted of protein biologics such as monoclonal antibodies, cytokines, and hormones.^2^ It is of note, however, that almost all approved protein therapies operate in extracellular environments, a restriction that is largely due to the inability of proteins to spontaneously enter cells. Indeed, cells have undergone billions of years of evolution to prevent the unassisted passage of such large molecular weight, hydrophilic macromolecules through their hydrophobic plasma membranes.

The potential of proteins as intracellular therapeutic agents has been underscored by the use of novel protein scaffolds that target the “undruggable” proteome,^3–5^ as well as the recent application of gene-editing proteins^6,7^ to treat human disease. Translating these promising technologies for clinically relevant applications requires the development of efficacious delivery methods capable of transporting proteins into the cytosol of human cells. While DNA or RNA encoding protein products can be delivered to cells via viral vectors or nanoparticles, these methods suffer from lack of temporal control, high immunogenicity, risk of genome integration, and unintended off-target effects *in vivo*.^8,9^ Viewed from this perspective, direct delivery of protein therapies into cells is a unique approach that overcomes some of the limitations and concerns associated with existing nucleic acid-based methods for treating different pathological conditions.

Over the years, several techniques have been pioneered for delivering proteins into the cytosol of cells, such as membrane disruption methods,^10,11^ chemical conjugation schemes (cell penetrating-peptides^12,13^ and hydrophobic “masking” compounds^14,15^), and carrier-mediated approaches (polymeric assemblies,^16,17^ virus-like particles,^18,19^ and inorganic nanostructures^20,21^). While these strategies achieved varying degrees of success for *in vitro* delivery, they suffer from key barriers that prevent their translation for clinical applications, including but not limited to low delivery efficiencies and instability in serum.^22^ The absence of FDA-approved protein delivery strategies highlights the considerable challenges associated with achieving successful intracellular delivery of protein therapies *in vivo*.

An emerging alternative strategy for protein delivery involves adapting methods that have already proven successful for the delivery of other biological drugs. In this vein, cationic lipids, best known for their ability to deliver nucleic acid cargos, have been recently explored as a promising platform for delivering proteins.^23–27^ Such cationic lipid carriers have enabled successful delivery of enzymes, CRISPR-Cas complexes, and inhibitory protein scaffolds with the potential for therapeutic use. However, most of these previous efforts required protein cargos to be genetically fused with anionic polypeptides or protein domains to promote electrostatic interactions with cationic lipids. Thus, the implementation of such strategies involves genetic manipulation to reengineer protein cargos with anionic tags, which can be time-consuming, may not always be tolerated by the cargo protein, and limits off-the-shelf proteins from being directly functionalized for delivery.

Owing to the clinical success of cationic lipids, particularly lipid nanoparticle (LNP) formulations that have garnered much attention during the COVID-19 pandemic,^28^ we were similarly interested in utilizing lipid-based carriers for delivery of protein therapeutics with an eye towards widespread generalizability and applicability. To this end, we explored a reversible bioconjugation strategy that endows proteins with an anionic “cloak” to facilitate electrostatic complexation with cationic lipids for intracellular delivery (**Fig. 1**). This is achieved through lysine-reactive activated carbonate compounds containing branched anionic sulfonate moieties that can efficiently remodel the surface charge of any given protein cargo. Further, by utilizing self-immolative disulfide chemistry, the compounds can be cleaved from the delivered proteins within the reducing environment of the cytosol, offering a traceless method of protein delivery. We establish proof-of-concept for this novel delivery approach and showcase its utility for the functional delivery of a variety of protein cargos, including a therapeutic enzyme and a full-length antibody, both *in vitro* and *in vivo*.

**Figure 1.**
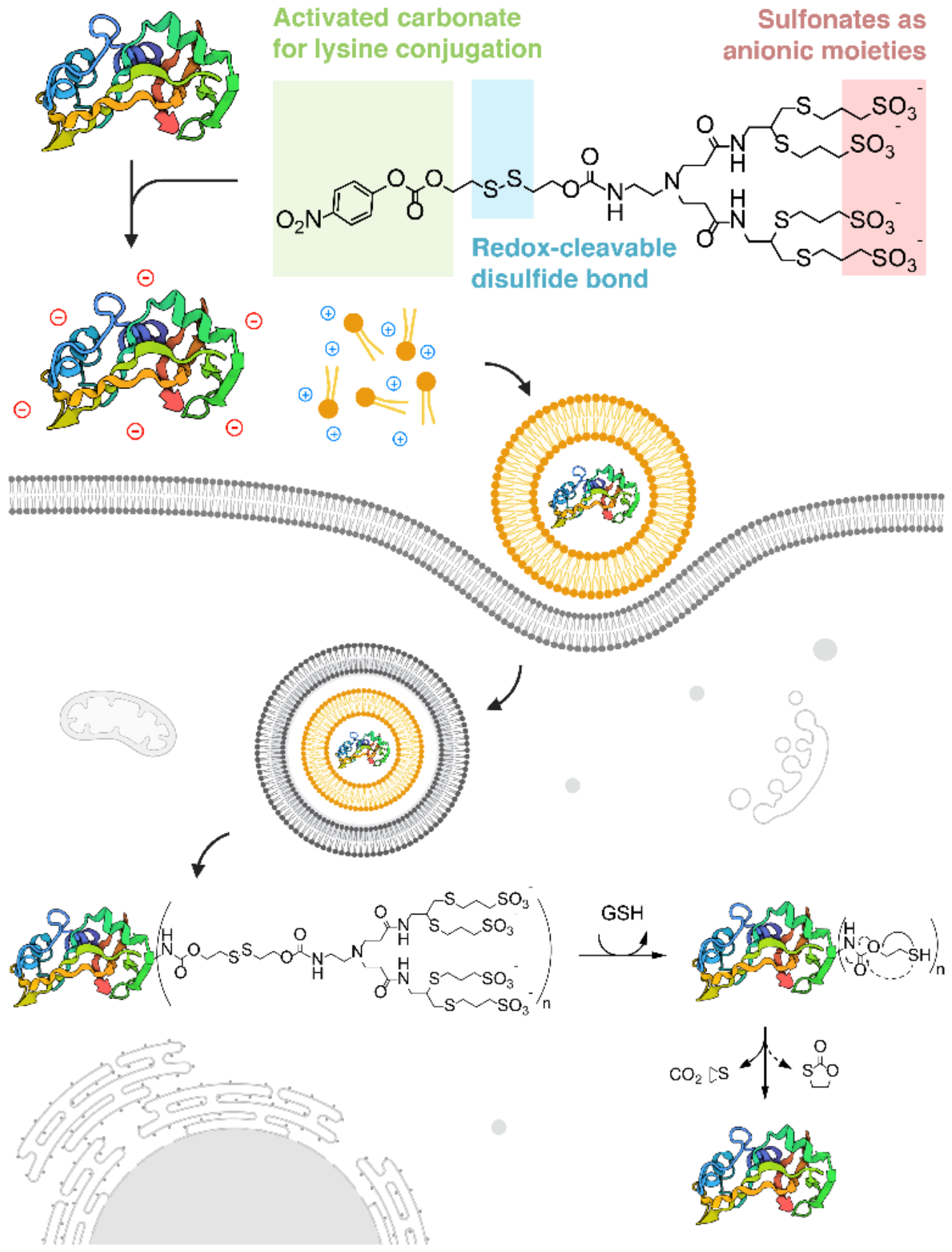
Schematic of the bioreversible anionic cloaking strategy. Chemical modification of surface-exposed lysines with sulfonated cloaking reagents enables complexation and subsequent delivery of protein cargos with cationic lipids. Following endocytic escape, the reagents are cleaved off via the presence of a self-immolative, redox-sensitive disulfide bond to tracelessly deliver the cargo protein.

## Results

### Modification of proteins with anionic cloaking reagents significantly remodels protein surface charge

We reasoned that the formation of an effective anionic cloak would require global surface charge remodeling to endow sufficient anionic character to a cargo protein. To test this notion, we investigated activated carbonate compounds to chemoselectively attach sulfonate groups to surface-exposed lysine residues on proteins of interest. This bioconjugation-based approach allows for global, nonspecific charge reversal of positively charged amines (lysine, histidine, or the N-terminal residue) to negatively charged sulfonate moieties via carbamate formation. Charge modification with sulfonates enables the formation of a strong anionic cloak due to their exceptionally low pK_a_ (pK_a_ ≈ -2). Additionally, incorporating a disulfide bond β-to the carbamate attachment enables redox-mediated cleavage and self-immolation to tracelessly regenerate the native protein within the reducing environment of the cytosol. The sulfonated *p*-nitrophenyl carbonate compounds containing disulfide linkers (**Fig. 2a**) were synthesized according to the chemical scheme in (**Supplementary Fig. S1**). Briefly, allylamine, diallylamine or propargylamine were acylated with acryloyl chloride, followed by a Michael addition with *N*-Boc-ethylenediamine. The anionic groups were installed by reacting the resulting allyl and propargyl compounds with 3-mercapto-1-propane sulfonate followed by Boc deprotection. SL2b was obtained from a thiyl radical-induced cyclization, wherein the nascent carbon radical formed led to a mixture of 5-*exo* and 6-*endo* cyclized products. Final compounds were purified via RP-HPLC and characterized via mass spectrometry and NMR (**Supplementary Fig. S9-S33)**.

**Figure 2.**
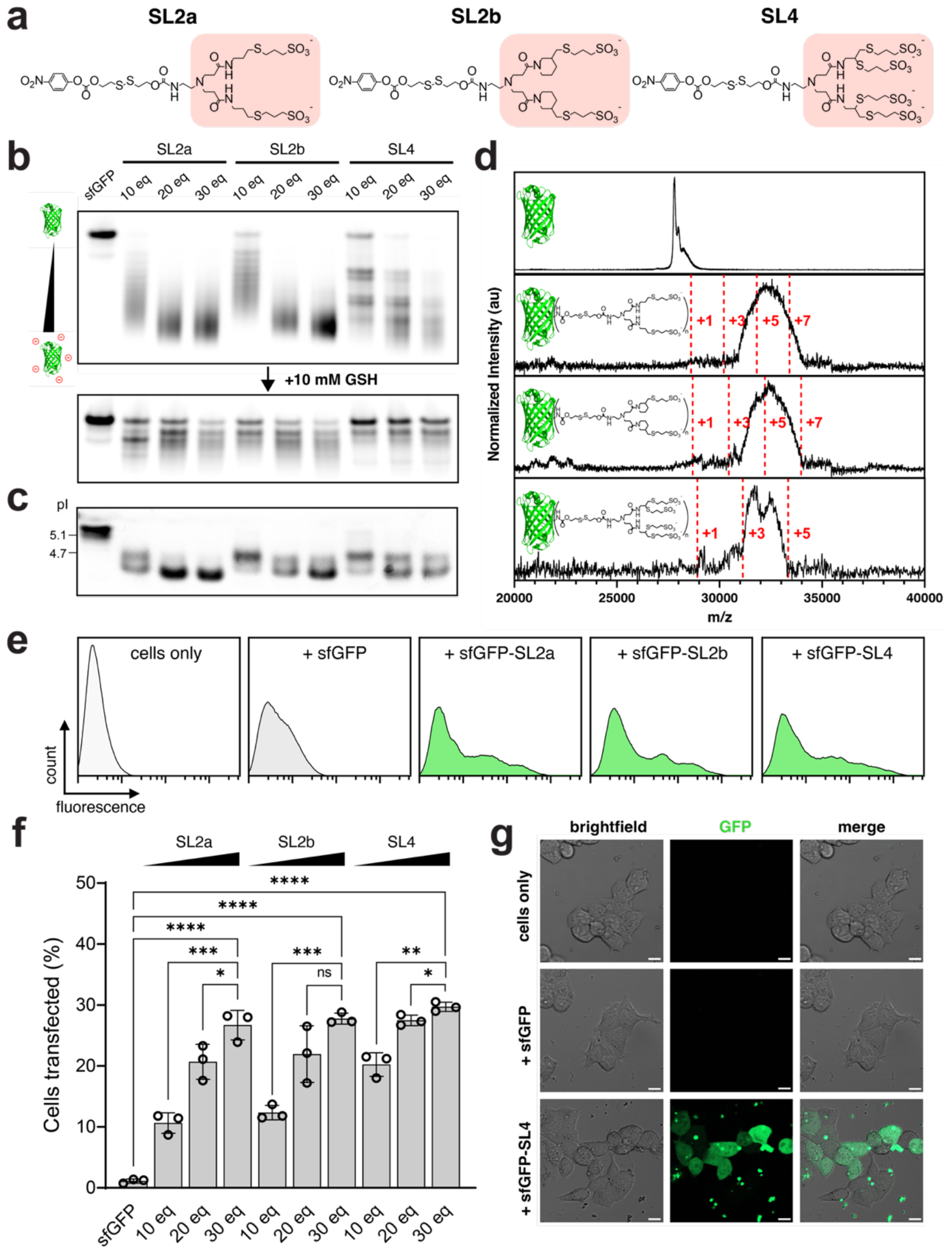
Conjugation of sfGFP with lysine-reactive sulfonated probes enables efficient delivery. Transfections of sfGFP complexed with Lipofectamine 2000 (LF2K) were performed at 500 nM into HEK293T cells for 6 hours. (a) Panel of sulfonated *p*-nitrophenyl carbonate compounds synthesized for this study. (b) Native gel electrophoresis of sfGFP samples conjugated to sulfonated compounds, before and after treatment with 10 mM GSH. (c) Isoelectric focusing gels of sfGFP conjugated to sulfonated compounds. (d) MALDI spectra of sfGFP samples conjugated to sulfonated compounds (modified with 30 molar equivalents). (e) Representative flow cytometry histograms of HEK293T cells transfected with sfGFP and sfGFP modified with 30 molar equivalents of each sulfonated compound, using LF2K. (f) Percent GFP-positive HEK293T cells following transfections of sfGFP and anionically-modified sfGFP (with molar equivalents of sulfonated compounds as indicated) using LF2K. (g) Representative confocal microscopy images of HEK293T cells transfected with sfGFP and sfGFP-SL4 (modified with 30 molar equivalents of SL4) using LF2K. Scale bar = 10 μm. All data are mean ± SD (*n* = 3 for flow cytometry). Statistical significance was determined by ordinary one-way ANOVA followed by Bonferroni correction for multiple comparisons (^*^*p* < 0.05, ^**^*p* < 0.01, ^***^*p* < 0.001, ^****^*p* < 0.0001).

To validate the anionic cloaking strategy, we employed superfolder green fluorescent protein (sfGFP) as a model protein to study the conjugation and delivery process. Successful anionic protein modification was confirmed by polyacrylamide gel electrophoresis run under native conditions (**Fig. 2b**). Addition of increasing molar equivalents of the three sulfonated compounds resulted in increasingly faster migration of the modified sfGFP through the gel towards the positive potential. It is worth noting that sfGFP retains its intrinsic fluorescence upon chemical modification, as evidenced by the in-gel fluorescence images. Furthermore, complete protein modification occurred when sfGFP was reacted with 30 molar equivalents of each of the sulfonated compounds. Incubation of the modified sfGFP samples with 10 mM GSH (corresponding to the approximate concentration in a reducing cytosolic environment^29^) resulted in convergence of the bands towards the unmodified sfGFP band, demonstrating successful disulfide cleavage and traceless recovery of the native protein. Anionic modification of sfGFP was further resolved through isoelectric focusing (IEF), which clearly showed a reduction in sfGFP isoelectric point (pI) to below 4.7 upon conjugation of the sulfonated compounds (**Fig. 2c**). Interestingly, the pI of sfGFP at full modification was approximately the same for all three sulfonated compounds, even though the compounds vary in overall hydrophobicity and valency of sulfonate groups. From MALDI-TOF-MS analysis, shifts in mass peaks arising from lysine-attached adducts revealed an average degree of conjugation ranging from 3 to 5 for the fully modified proteins (**Fig. 2d**). The resulting bioconjugation thus corresponds to an estimated decrease in the theoretical net charge of sfGFP from approximately -2 to a range between -11 to -27 under physiological conditions.

### Anionic cloaking enables intracellular protein delivery with commercial cationic lipid reagents

Having established efficient anionic bioconjugation, we next investigated the delivery of anionically-cloaked sfGFP into cells using Lipofectamine 2000 (LF2K), a commercial cationic lipid reagent routinely employed for *in vitro* transfection of nucleic acids. Flow cytometry experiments with 500 nM of anionically-cloaked sfGFP complexed with LF2K revealed elevated intracellular fluorescence in HEK293T cells (**Fig. 2e**), indicative of successful protein internalization. Conversely, cells treated with native, unmodified sfGFP complexed with LF2K exhibited no measurable delivery. Furthermore, fluorescent signals in cells were observed with sfGFP concentrations as low as 50 nM (**Supplementary Fig. S2**). Efficiency of LF2K-mediated sfGFP delivery (quantified as percent GFP-positive cells) into cells increased with increasing amounts of sulfonate modification (**Fig. 2f**), suggesting that the degree of anionic protein modification may correlate with efficiency of electrostatic complexation with cationic lipids and, ultimately, delivery efficiency. However, maximal delivery efficiency with LF2K was capped at 30% for sfGFP that was reacted with 30 molar equivalents of all three sulfonated compounds. Confocal microscopy images corroborated the flow cytometry results and confirmed that protein internalization within cells only occurred when LF2K was complexed with anionically-cloaked sfGFP (**Fig. 2g**). Taken together, these results demonstrate that a ∼30 kDa globular protein can undergo anionic surface charge remodeling to enable electrostatic complexation and intracellular delivery with off-the-shelf cationic lipid reagents.

### Anionically-cloaked sfGFP formulated with LNPs is robustly internalized

Having established proof-of-principle of our protein delivery approach, we next sought to adapt our strategy for therapeutically relevant applications by utilizing clinically validated LNP formulations. Traditional LNP formulations consist of four lipid components – “ionizable” tertiary-amine containing lipids, zwitterionic phospholipids, cholesterol, and poly(ethylene) glycol (PEGylated) lipids – that are mixed at precise molar ratios to yield structured, homogenous nanoparticles. Essential to LNP formation with nucleic acids is the charge state of the ionizable lipid, which is modulated by the pH of the formulation mixture. In particular, ionizable lipids with pK_a_ ≈ 6.5 are able to (i) form electrostatic complexes with nucleic acids in acidic environments (e.g., buffers at pH 3), wherein the tertiary amines are protonated, and (ii) transition to an uncharged state at the physiological pH of 7.4. This feature is advantageous for minimizing off-target cytotoxicity during circulation.

Here, we formulated LNPs with anionically-cloaked sfGFP (modified with 30 molar equivalents of SL4) using the “gold standard” ionizable lipid DLin-MC3-DMA (MC3) utilized in the FDA-approved siRNA-based drug Onpattro.^30^ LNPs of varying lipid amounts (2-10 wt/wt, MC3/sfGFP) were formed using a traditional four component system comprised of MC3, distearoylphosphatidylcholine (DSPC), cholesterol, and distearoyl-*rac*-glycerol-methoxypoly(ethylene) glycol (DSG-PEG) (50/10/38.5/1.5 mol/mol). One concern is that traditional LNP formulations involve rapid mixing of an ethanolic lipid solution with an acidic aqueous buffer containing the nucleic acid. The application of such a harsh method to protein cargos could induce unintended disruption of protein structure and folding during the formulation process. Indeed, sfGFP fluorescence was quenched when the protein was placed in citrate buffer at pH 3 but retained its fluorescence at higher pH ranges (**Supplementary Fig. S3**). Shifting to higher formulation pH, however, could result in decreased populations of protonated ionizable lipids. To balance these competing effects, we hypothesized that the introduction of an auxiliary cationic lipid to the conventional four component LNP system would facilitate the electrostatic-driven assembly of LNPs with anionically-cloaked proteins in protein-friendly neutral pH buffers. To test this hypothesis, we generated an additional formulation in which the above four component mixture was supplemented with 10 mol% of 1,2-dioleoyl-3-trimethylammonium-propane (DOTAP), a cationic lipid consisting of a permanently charged quaternary ammonium. Dynamic light scattering measurements (DLS) revealed successful formation of nanoparticles ranging between 200-300 nm in size with low polydispersity across all formulations made at both pH 5 and pH 7.4 (**Supplementary Table S1**). Zeta potential measurements of the nanoparticles ranged from 0 and -5, signifying their overall neutral surface charge (**Supplementary Table S1**). Interestingly, encapsulation efficiency of sulfonate-cloaked sfGFP was markedly higher when formulated with LNPs supplemented with DOTAP in both pH 5 and pH 7.4 buffers, suggesting that DOTAP may play a crucial role in increasing protein encapsulation in LNPs (**Supplementary Fig. S4a**,**b**).

To examine whether these LNPs were capable of transporting sfGFP into cells, we transfected HEK293T cells with the above formulations and observed a pronounced shift in intracellular fluorescence for anionically-cloaked sfGFP complexed with LNPs that were supplemented with 10 mol% DOTAP (**Fig. 3a**). In stark contrast, there was little evidence of protein delivery for MC3 LNPs formulated with unmodified sfGFP or LNPs formulated using the traditional four component lipid system. The extent of protein delivery was particularly impressive when analyzing the delivery efficiencies, with nearly 90% of cells transfected with MC3 LNPs formulated with sfGFP-SL4 (**Fig. 3b**). Intracellular fluorescence was discernable in a dose-dependent manner, ranging from concentrations of sfGFP-SL4 at 250 nM down to 10 nM (**Fig. 3c**). Delivery of sfGFP-SL4 was also achieved using formulations of ALC-0315 and SM-102 - the ionizable lipids used in the SARS-CoV-2 mRNA vaccines from Pfizer/BioNTech and Moderna,^28^ respectively - but only with formulations supplemented with DOTAP (**Fig. 3d**). These results illustrate the adaptability of the anionic cloaking strategy across various LNPs comprised of different ionizable lipid architectures but with the caveat that supplementation with an additional cationic lipid is necessary for LNP-mediated protein delivery. Furthermore, LNP-mediated protein delivery is nontoxic, with most cells exhibiting viabilities above 90% following treatment (**Fig. 3e**). This promising biocompatibility bodes well for potential *in vivo* applications. We next investigated the effect of charge type on the anionic-cloaking mechanism by modifying sfGFP with *p*-nitrophenyl carbonate compounds containing carboxylate moieties (see **Supplementary Fig. S1** for details on synthesis). In principle, surface charge modification with carboxylates, which possess a much higher pK_a_ (pK_a_ ≈ 5) compared to sulfonates, would result in a weaker anionic cloak at a formulation pH near or above this pKa and, subsequently, less efficient complexation with ionizable lipids. This weaker anionic cloak was indeed evidenced by a reduced shift in the pI of sfGFP when it was cloaked with 30 molar equivalents of the carboxylated compound, CL4, compared to that of the sulfonated compound, SL4 (**Supplementary Fig. S5**). As a result, encapsulation efficiency of CL4-cloaked sfGFP (sfGFP-CL4) within LNPs was notably reduced compared to SL4-cloaked sfGFP (sfGFP-SL4) (**Supplementary Fig. S4c**,**d**). Consistent with the previously observed trend, supplementing the LNP formulation with DOTAP, both at pH 5 and pH 7.4, led to improved encapsulation efficiency for sfGFP-CL4 (**Supplementary Fig. S4c**,**d**). This result further emphasizes the vital contribution of DOTAP in enhancing protein encapsulation with cationic lipids. Following transfection of HEK293T cells with MC3 LNPs formulated with sfGFP-CL4, protein delivery was observed, albeit at a level lower than with sfGFP-SL4 (**Fig. 3a,b**).

**Figure 3.**
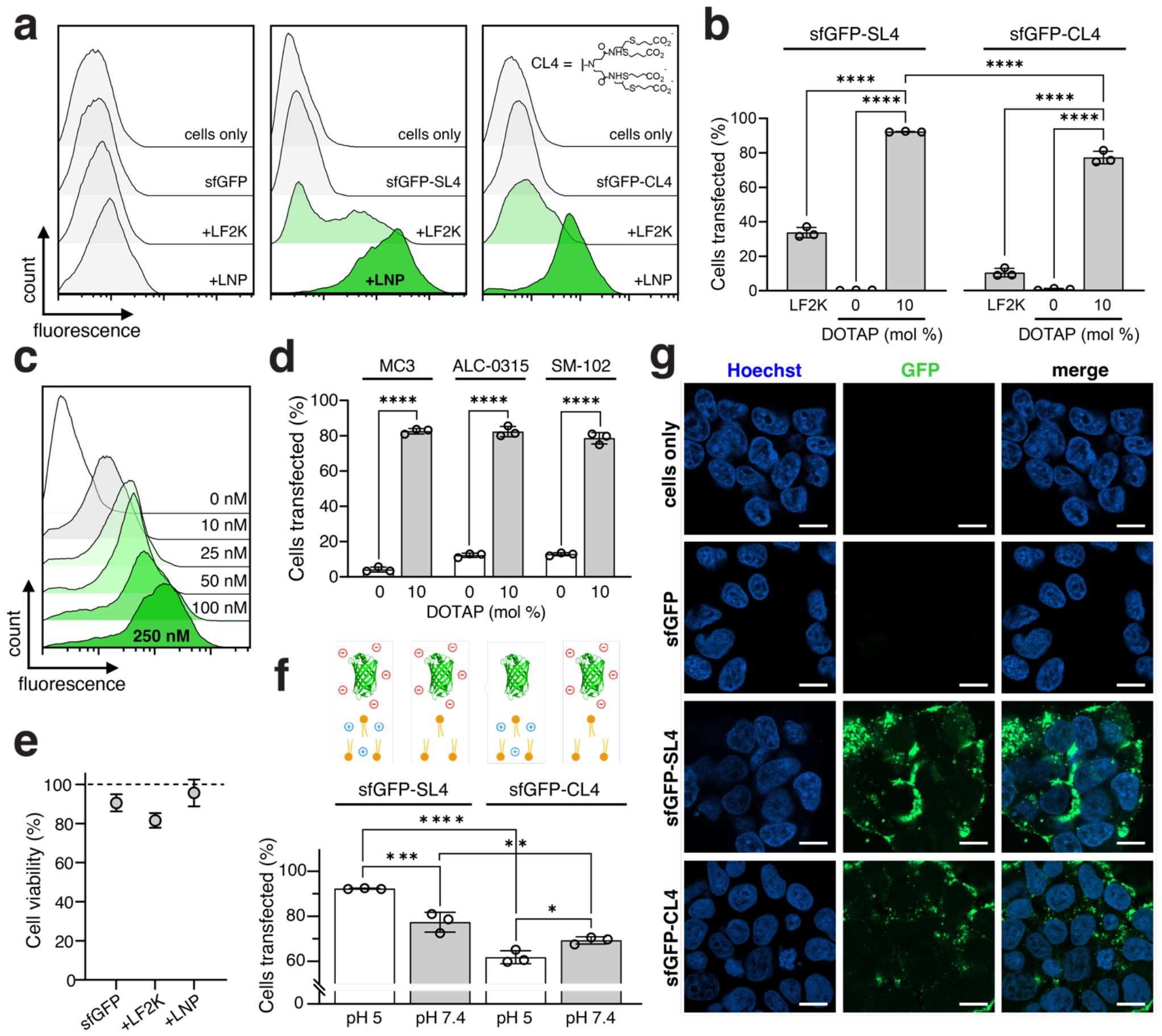
Delivery of anionically-cloaked sfGFP with MC3 LNPs. Transfections of sfGFP using MC3 LNPs were performed at 250 nM into HEK293T cells for 6 hours. Unless otherwise stated, data shown are for sfGFP cloaked with 30 molar equivalents of SL4 or CL4 and for MC3 LNPs (10 wt/wt, MC3/sfGFP) supplemented with 10 mol% DOTAP and formulated in pH 5 citrate buffer. (a) Representative flow cytometry histograms of HEK293T cells transfected with sfGFP and sfGFP cloaked with SL4 or CL4, using MC3 LNPs. For comparison, HEK293T cells were transfected with 500 nM of the same sfGFP proteins complexed with LF2K. (b) Percent GFP-positive HEK293T cells following transfections of sfGFP-SL4 and sfGFP-CL4 using LF2K and MC3 LNPs. (c) Representative flow cytometry histograms of HEK293T cells transfected with 10–250 nM of sfGFP-SL4 using MC3 LNPs. (d) Percent GFP-positive HEK293T cells following transfections of sfGFP-SL4 using MC3, ALC-0315, and SM-102 LNPs. LNPs (10 wt/wt, ionizable lipid/sfGFP) were supplemented with 10 mol% DOTAP and formulated in pH 5 citrate buffer. (e) Viability of HEK293T cells following transfections of sfGFP alone and sfGFP-SL4 using LF2K and MC3 LNPs, as measured by MTS assay. (f) Percent GFP-positive HEK293T cells following transfections of sfGFP-SL4 and sfGFP-CL4 using MC3 LNPs formulated in citrate buffers at pH 5 and pH 7.4. (g) Representative confocal microscopy images of HEK293T cells transfected with sfGFP, sfGFP-SL4, and sfGFP-CL4 using MC3 LNPs. Scale bar = 10 μm. All data are mean ± SD (*n* = 3 for flow cytometry; *n* = 4 for MTS). Statistical significance was determined by unpaired *t-*tests followed by Bonferroni-Dunn correction for multiple comparisons (\**p* < 0.05, ^**^*p* < 0.01, ^***^*p* < 0.001, ^****^*p* < 0.0001).

Comparing delivery between sulfonate versus carboxylate-cloaked sfGFP, it was evident that surface charge modification with sulfonates led to enhanced delivery using both LF2K and MC3 LNPs (**Fig. 3b**), owing to the increased anionic character of the modified proteins. The differences in delivery between the two anionic cloaks is supported by confocal microscopy images, where a more prominent GFP signal was observed in cells treated with sfGFP-SL4 compared to sfGFP-CL4 (**Fig. 3g**).

Specifically, delivery was higher for MC3 LNPs formulated with sfGFP-SL4 at both pH 5 and Ph 7.4 of mixing compared to those formulated with sfGFP-CL4 (**Fig. 3f**). Interestingly, the highest protein delivery was achieved at pH 5 of mixing with sfGFP-SL4, whereas delivery of sfGFP-CL4 appeared roughly similar at both pH 5 and pH 7.4. The trends in delivery efficiency tended to correlate with sfGFP encapsulation efficiency of formulated LNPs, with sfGFP-SL4 achieving >70% encapsulation efficiency at pH 5 and ∼40% at pH 7.4, whereas encapsulation efficiency of sfGFP-CL4 was ∼30% when formulated with LNPs in both pH 5 and pH 7.4 buffers (**Supplementary Fig. S4a-d**).

### LNP-mediated delivery of cloaked RNase A induces potent cytotoxity

To expand the scope of our delivery platform, we next investigated the functional delivery of ribonuclease A (RNase A), a 13.7-kDa endonuclease that cleaves single-stranded RNAs. High levels of RNase A inside cells induces cytotoxic effects,^31^ which has motivated efforts to deliver RNase A using polymeric- and lipid-based materials^32–34^ for potential anticancer applications. Here, we sought to apply our cloaking strategy to deliver RNase A with LNPs into different cancer cell lines and use cytotoxicity as a phenotypic surrogate to evaluate functional delivery (**Fig. 4a**).

**Figure 4.**
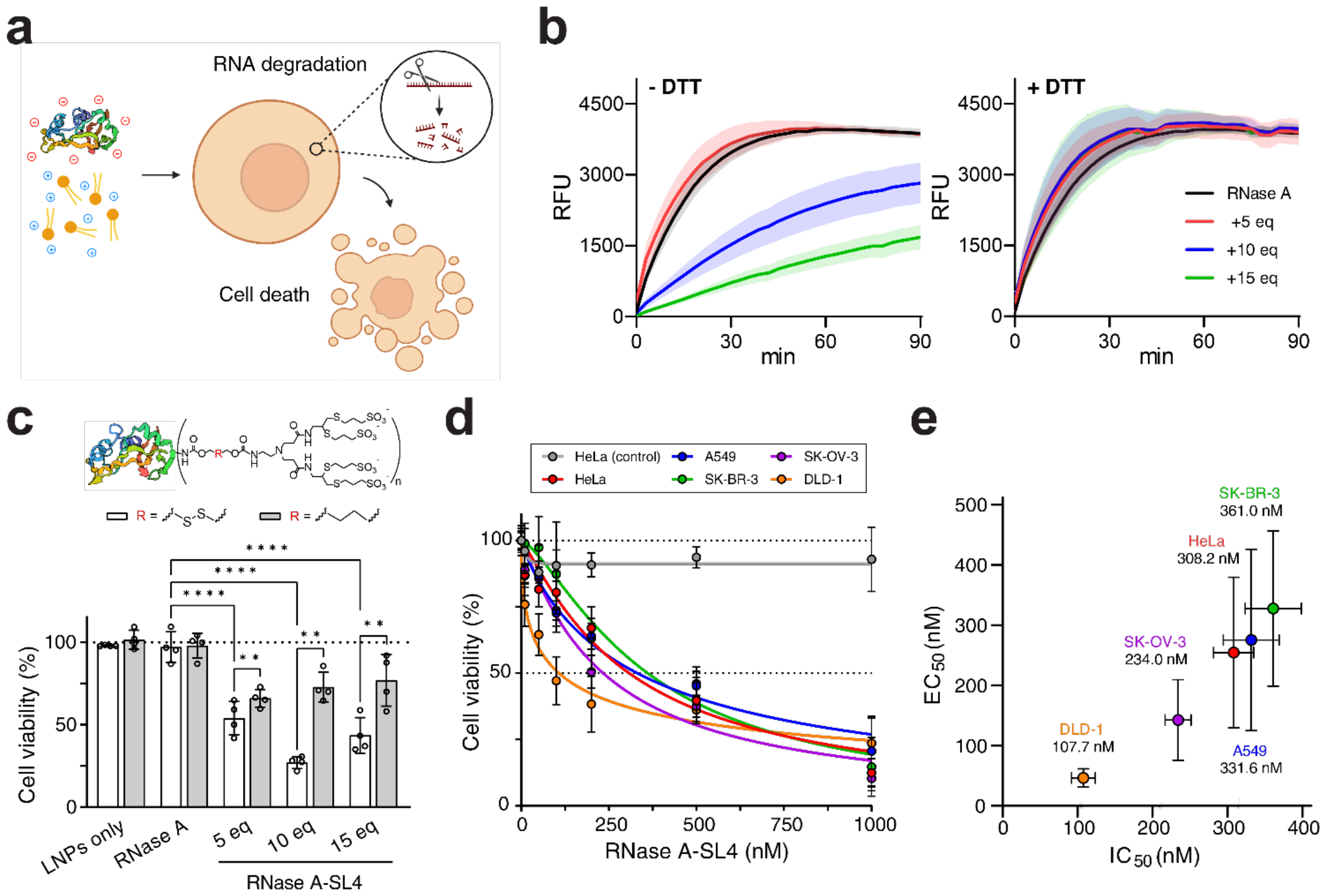
Delivery of anionically-cloaked RNase A with MC3 LNPs. Data shown are for MC3 LNPs (10 wt/wt, MC3/RNase A) supplemented with 10 mol% DOTAP and formulated in pH 5 citrate buffer. Unless otherwise stated, all RNase A transfections were performed for 48 hours. (a) Schematic depicting RNase A delivery strategy. Following delivery, RNase A will induce degradation of intracellular RNA, leading to cell death. (b) Ribonuclease activity of native RNase A and RNase A-SL4. Activity assays were repeated for RNase A samples incubated overnight with 10 mM DTT. (c) Viability of HEK293T cells following 500 nM transfections of RNase A-SL4 using MC3 LNPs, as measured by MTS assay. RNase A was modified with 5-15 molar equivalents of SL4 either containing redox-cleavable disulfide bonds or non-cleavable butyl linker. (d) Viability of cancer cell lines following transfections of RNase A-SL4 from 10 nM – 1000 nM using MC3 LNPs, as measured by MTS assay. RNase A was modified with 10 molar equivalents of SL4. HeLa cells treated with blank MC3 LNPs (gray line) at equivalent lipid amounts served as a negative control. (e) Half-maximal effective concentration of delivery, EC_50_, vs IC_50_ values for cells transfected with RNase A-SL4 using MC3 LNPs. RNase A was modified with 10 molar equivalents of SL4. Calculated IC_50_ values are shown under each cell line. EC_50_ values calculated by transfecting cells with 10–500 nM of fluorescein-labeled RNase A-SL4 using MC3 LNPs for 6 hours and quantifying percent positive fluorescein-RNase A-SL4 cells using flow cytometry. All data are mean ± SD (*n* = 3 for flow cytometry; *n* = 4 for MTS; *n* = 4 for ribonuclease assay). Statistical significance was determined by two-way ANOVA followed by Bonferroni correction for multiple comparisons ^(*^*p* < 0.05, ^**^*p* < 0.01, ^***^*p* < 0.001, ^****^*p* < 0.0001).

We assessed conjugation and anionic modification of RNase A with SL4. RNase A is a highly basic protein (pI ≈ 8.5) and was readily conjugated with as little as 5 molar equivalents of SL4, resulting in a pI below 5 (**Supplementary Fig. S6a**). MALDI-TOF-MS analysis confirmed the attachment of 3 to 5 sulfonated compounds to RNase A modified with 5 to 15 molar equivalents of SL4 (**Supplementary Fig. S6b**). Circular dichroism (CD) spectra revealed no discernable changes in the secondary structure of RNase A after cloaking with SL4 and after DTT-mediated reduction of cloaked RNase A (**Supplementary Fig. S6c**).

Imperative for functional protein delivery is the ability of a protein cargo to retain its biological activity upon chemical modification. We therefore evaluated the activity of RNase A cloaked with SL4 using a standard ribonuclease assay kit. Modification of RNase A with increasing molar equivalents of SL4 reduced nuclease activity, particularly when reacted with 10 or higher molar equivalents of SL4 (**Fig. 4b**). However, incubation of the cloaked RNase A samples with 10 mM DTT prior to measuring activity resulted in complete recovery of enzymatic activity back to that of the unmodified enzyme, indicating that reductive cleavage of the sulfonated compounds can restore native protein function. This is further corroborated by recovery of the basic RNase A band in the IEF gel upon incubation with 10 mM GSH, demonstrating successful disulfide linker cleavage and traceless recovery of the native protein (**Supplementary Fig. S6a**). Interestingly, co-incubation of cloaked RNase A with 10 mM of either GSH or DTT resulted in gradual recovery of enzyme activity over the course of 6 hours and highlights that cleavage kinetics may play an important role in recovering protein function (**Supplementary Fig. S6d**).

To optimize RNase A formulations for cellular cytotoxicity, we evaluated various MC3 LNPs (supplemented with 10 mol% DOTAP) for delivery of anionically-cloaked RNase A into HEK293T cells. Transfections of RNase A cloaked with 5 to 15 molar equivalents of SL4 and formulated into LNPs in pH 5 buffer with varying amounts of lipids (1-10 wt/wt, MC3/RNase A) were all observed to reduce viability of HEK293T cells (**Supplementary Fig. S6e**). Maximal reduction in cell viability of nearly 70% was achieved with RNase A-SL4 modified with 10 molar equivalents and formulated with 10 wt/wt, MC3/RNase A (**Fig. 4c** and **Supplementary Fig. S6e**). To evaluate the importance of intracellular disulfide linker cleavage for recovery of enzymatic activity, transfections were also performed with RNase A modified with non-redox-cleavable variants of SL4, which led to a less significant reduction in cell viability compared to that of RNase A modified with cleavable compounds (**Fig. 4c**). The diminished activity of RNase A modified by non-cleavable SL4 was further confirmed from ribonuclease activity assays that demonstrated no recovery in RNase A activity following incubation of cloaked RNase A with 10 mM DTT (**Supplementary Fig. S6g**).

We next explored cytotoxicity against a wider range of clinically relevant cancer cell lines, including A549 (lung), DLD-1 (colorectal), HeLa (cervical), SK-BR-3 (breast), and SK-OV-3 (ovarian), that vary in size, gene expression profiles, signaling pathways, DNA repair capacity, and cell cycle regulation. Treatment with RNase A modified with 10 molar equivalents of SL4 and formulated with 10 wt/wt, MC3/RNase A resulted in a potent dose-dependent reduction in the viability of all tested cancer cells (**Fig. 4d**), with calculated IC_50_ values below 400 nM in each case (**Fig. 4e**). To better understand the differential responses to RNase A treatment, we performed transfections of fluorescein-labeled RNase A-SL4 to quantify the extent of uptake into cells. The half-maximal effective concentration of uptake, EC_50_, correlated well with the calculated IC_50_ values for all cancer cells, indicating that the degree of cytotoxicity induced depends on the amount of RNase A delivered (**Fig. 4e**). Taken together, these results demonstrate that our anionic cloaking method enables efficient delivery of RNase A into a wide variety of cancer cell lines for cancer therapy applications.

### Delivery of inhibitory antibodies downregulates β-catenin activity

Immunoglobulin (IgG) antibodies, which possess high affinity and specificity towards their targets, are being increasingly explored for inhibition of intracellular signaling pathways and “undruggable” protein targets.^17,25,35–37^ To this end, we first investigated cloaking and delivery of a fluorescently labeled mouse anti-human IgG to optimize cellular internalization of this large, complex protein cargo. Conjugation with at least 30 molar equivalents of SL4 reduced the pI of the IgG to ∼5 (**Supplementary Fig. S7a**). Reacting beyond 30 molar equivalents of SL4 had no significant additional impact on pI. Anionically-cloaked IgG was then formulated into LNPs in pH 5 buffer at 2 wt/wt, MC3/antibody and transfected in HEK293T cells. We kept the lipid/protein weight ratio low due to the large molecular weight of the IgG. Flow cytometry analysis revealed that mouse anti-rabbit IgG cloaked with 15-60 molar equivalents of SL4 exhibited 60-80% delivery efficiency into cells (**Supplementary Fig. S7b**). Notably and consistent with conjugation experiments, delivery efficiency reached a plateau for mouse anti-rabbit IgG modified with over 30 molar equivalents of SL4. We reasoned that cloaking of IgG with 30 molar equivalents of SL4 was optimal given that protein pI and efficiency of delivery with LNPs do not increase significantly with further modification. Importantly, free IgG antibody in solution and uncloaked IgG antibodies formulated with MC3 LNPs did not internalize into cells (**Supplementary Fig. S7c**), reaffirming that the anionic cloaking mechanism is necessary for robust intracellular antibody delivery with LNPs.

We next investigated the delivery of a murine monoclonal IgG antibody specific for β-catenin using our anionic cloaking strategy and LNPs. We selected the transcription factor β-catenin as a target protein for the delivery of inhibitory antibodies because it plays a pivotal role in oncogenic Wnt transduction pathways.^38^ In Wnt-driven cancer pathogenesis, aberrantly stabilized β-catenin accumulates in the cytosol, translocates to the nucleus, and interacts with TCF/LEF transcription complex to drive the expression of oncogenes including *c-Myc* and *cyclin D1*.^38^ To date, there have been no approved therapies against β-catenin, although a few are currently undergoing clinical trials.^39^ We hypothesized that anionically-cloaked anti-β-catenin IgG antibodies complexed with MC3 LNPs could be delivered to the cytosol where they would bind stabilized β-catenin and prevent it from entering the nucleus, thereby inhibiting its transcriptional activity (**Fig. 5a**).

**Figure 5.**
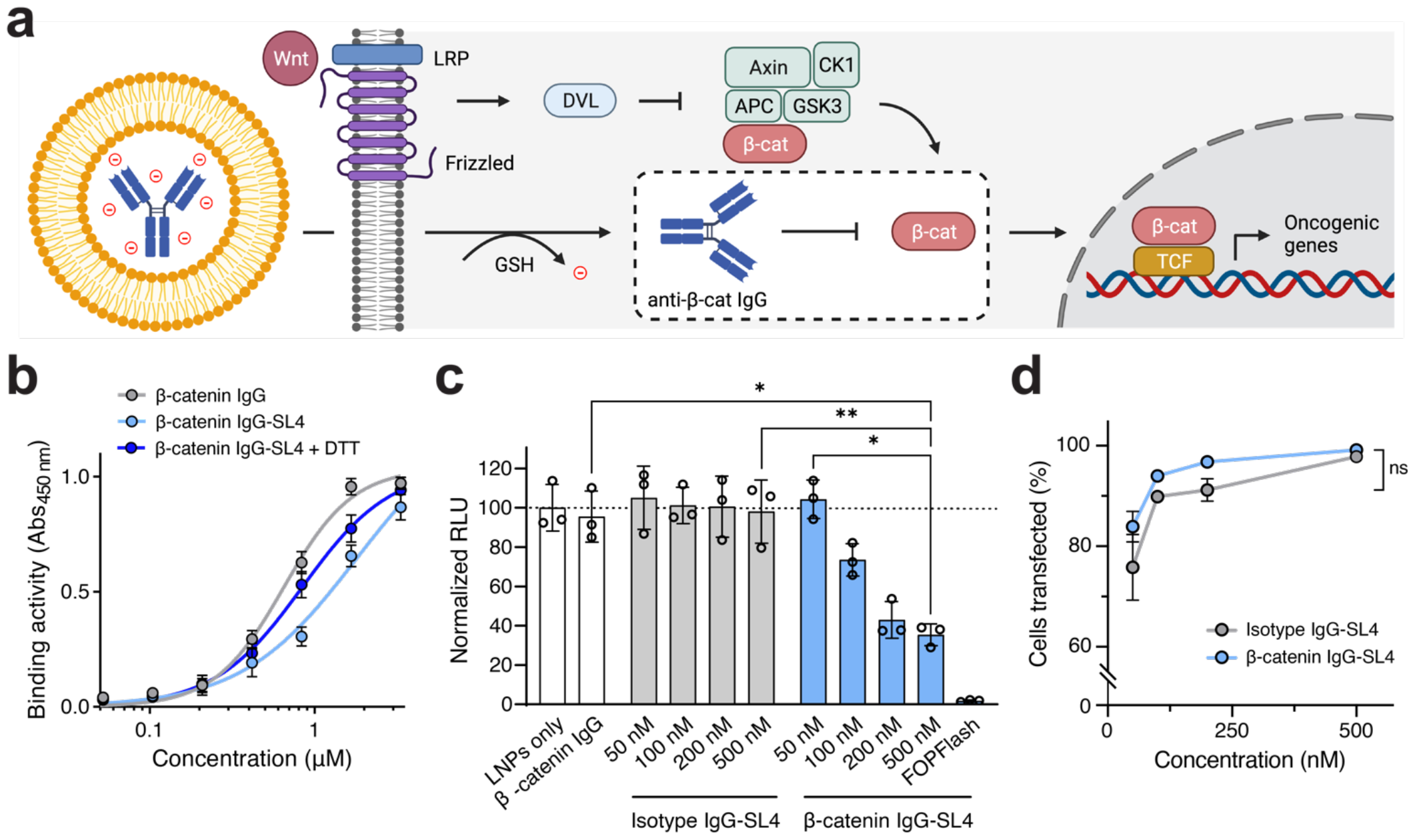
Delivery of anionically-cloaked anti-β-catenin antibody with MC3 LNPs. Data shown are for IgGs cloaked with 30 molar eq. of SL4 and for MC3 LNPs (MC3/IgG, 2 wt/wt) supplemented with 10 mol % DOTAP and formulated in pH 5 citrate buffer. (a) Schematic of IgG delivery strategy. Binding of β-catenin by anti-β-catenin IgG will prevent association of β-catenin with TCF and inhibit expression of Wnt-driven genes. (b) Binding activity of anti-β-catenin IgG to immobilized β-catenin as determined by ELISA in the presence or absence of DTT. IgGs were cloaked with 30 molar equivalents of SL4. (c) Knockdown of TCF-driven TOPFlash luciferase activity following transfection of DLD-1 cells with 50–500 nM anti-β-catenin IgG-SL4 and isotype IgG-SL4 using MC3 LNPs. DLD-1 cells transfected with FOPFlash plasmid, which contains mutated TCF sites upstream of luciferase expression cassette, served as a negative control. Transfections were performed for 24 hours. (d) Percent of fluorescein-IgG positive DLD-1 cells following transfections of 50-500 nM fluorescein-labeled anti-β-catenin IgG-SL4 and isotype IgG-SL4 using MC3 LNPs for 6 hours. IgGs were cloaked with 30 molar equivalents of SL4. All data are mean ± SD (*n* = 3 for flow cytometry; *n* = 3 for ELISA; *n* = 3 for TOPFlash assay). Statistical significance was determined by two-way ANOVA followed by Bonferroni correction for multiple comparisons ^(*\**^*p* < 0.05, ^*\*\**^*p* < 0.01, ^*\*\*\**^*p* < 0.001, ^*\*\*\*\**^*p* < 0.0001).

To test this hypothesis, we subjected the anti-β-catenin IgG to our conjugation strategy. MALDI-MS analysis confirmed conjugation and revealed an average degree of labeling between 3-5 when reacted with 30 molar equivalents of SL4 (**Supplementary Fig. S7d**). The ability of the antibody to bind β-catenin following SL4 conjugation was assessed via quantitative ELISA. While cloaking of the anti-β-catenin IgG with SL4 reduced binding activity to β-catenin, strong binding was largely restored following reduction of anti-β-catenin IgG-SL4 in the presence of 10 mM DTT (**Fig. 5b**), suggesting that binding of intracellular β-catenin is possible after cleavage of the anionic cloak following cytosolic delivery.

To test functional β-catenin inhibition, we leveraged the TOPFlash assay, a β-catenin-responsive plasmid reporter consisting of TCF binding sites placed upstream of a luciferase expression casette.^40^ As expected, constitutively Wnt-active DLD-1 colorectal cancer cells treated with LNPs only or anti-β-catenin IgG alone exhibited a strong TOPFlash signal (**Fig. 5c**), indicative of strong β-catenin-mediated transcriptional activity. In contrast, DLD-1 cells treated with 50–500 nM of anionically-cloaked anti-β-catenin IgG, but not cloaked isotype control IgG, delivered with MC3 LNPs exhibited a substantial dose-dependent reduction of the TOPFlash signal (**Fig. 5c**). It should be noted that delivery with as little as 200 nM of anti-β-catenin IgG-SL4 resulted in >60% reduction of β-catenin transcriptional activity. Using a fluoresein-labeled anti-β-catenin IgG-SL4 complexed with MC3 LNPs, we found that uptake into DLD-1 cells was highly efficient with ∼100% transfection efficiency at 500 nM treatments as determined by flow cytometry (**Fig. 5d**). Nearly identical transfection efficiency was observed for a fluorescein-labeled isotype control IgG-SL4 across the tested concentration ranges, suggesting that reduction in the TOPFlash signal resulted from specific binding and sequestration of β-catenin following delivery of the anti-β-catenin antibody. Overall, these findings demonstrate the feasibility of employing a commercially available antibody for intracellular cell signaling modulation, thereby paving the way for repurposing other off-the-shelf antibodies for various biological and therapeutic applications.

### LNPs distribute anionically-cloaked mCherry to major organs *in vivo*

To investigate the potential of our approach for *in vivo* applications, we investigated the biodistribution of anionically-cloaked mCherry protein with MC3 LNPs following systemic administration in mice. For *in vivo* studies, we first optimized LNP formulations with cloaked mCherry (10 wt/wt, total lipids/mCherry) by varying the amount of DOTAP (10-30 mol%) and PEG-DMG-2000 (1.5-4.5 mol%) and assessing each of these formulations for their average particle size, stability in mouse serum, transfection efficiency, and cellular cytotoxicity. Increasing both DOTAP and PEG-DMG-2000 resulted in particles with decreasing size, and additional DOTAP generally aided in serum stability and transfection efficiency (**Fig. 6a**). Increasing the amount of PEG-DMG-2000, however, had an adverse effect on cellular uptake of protein (**Fig. 6a**). Screening nine different formulations identified one (30 mol% DOTAP and 3 mol% PEG-DMG-2000) that was optimal in all four categories and thus was selected for use in biodistribution experiments.

**Figure 6.**
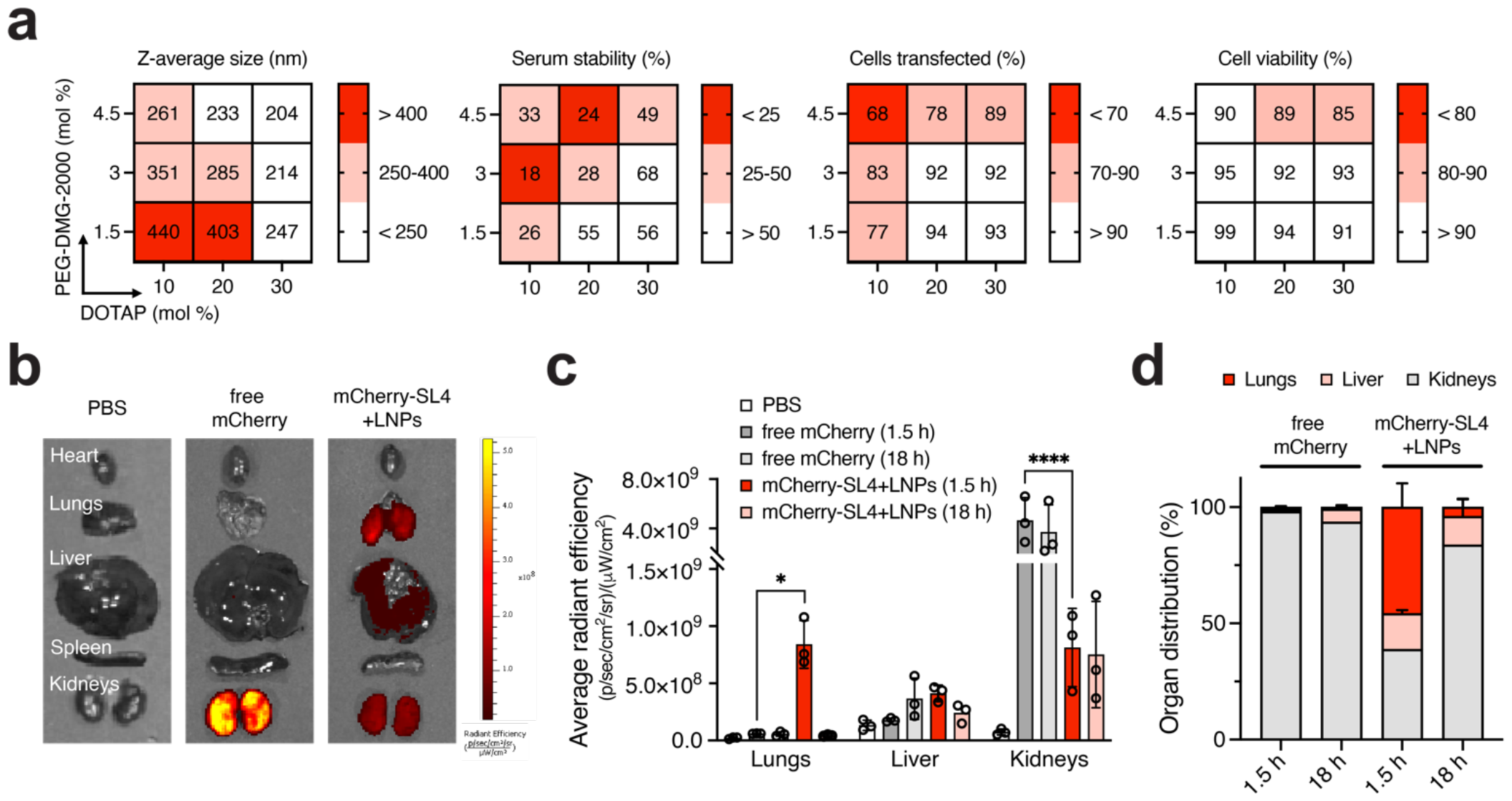
*In vivo* biodistribution of anionically-cloaked mCherry delivered with MC3 LNPs. Data shown are for mCherry cloaked with 30 molar equivalents of SL4 and for MC3 LNPs formulated in pH 5 citrate buffer. (a) Heat map of LNP formulation optimization of anionically-cloaked mCherry. MC3 LNPs were formulated with mCherry-SL4 using varying amounts of PEG-DMG-2000 and DOTAP (20 wt/wt, total lipids/mCherry). Average size distribution, serum stability, transfection efficiency into HEK293T cells, and cellular cytotoxicity were measured for the nine formulations. (b) Representative *ex vivo* fluorescent images of harvested organs following tail vein injection of SKH1 mice with PBS, free mCherry, and mCherry-SL4 formulated in MC3 LNPs. MC3 LNPs were formulated with 3 mol% PEG-DMG-2000 and 30 mol% DOTAP. Doses were 1 mg/kg of total protein. Images shown are for 1.5 h post-injection. (c) Quantified average radiant efficiency of *ex vivo* fluorescent images of harvested lungs, liver, and kidneys from SKH1 mice. (d) Average organ distribution of mCherry from harvest lungs, liver and kidneys from SKH1 mice. Percentage distribution was calculated by subtracting average PBS radiance from samples. Unless otherwise noted, all data are mean ± SD (*n* = 3 for flow cytometry; *n* = 4 for MTS; *n* = 3 for mice injections). Statistical significance was determined by two-way ANOVA followed by Bonferroni correction for multiple comparisons ^(*^*p* < 0.05, ^**^*p* < 0.01, ^***^*p* < 0.001, ^****^*p* < 0.0001).

Following tail-vein injection of mice with anionically-cloaked mCherry (modified with 30 molar equivalents of SL4; **Supplementary Fig. S8a**) encapsulated in MC3 LNPs, we observed biodistribution primarily to kidneys, liver, and lungs with ∼45% of the mCherry dose localizing to the lungs in 1.5 hours (**Fig. 6b-d** and **Supplementary Fig. S8b**). This finding is consistent with recent reports demonstrating that LNPs modified to be cationic through supplementation with excipient cationic lipids can shift LNP tissue tropism from liver to lungs due to the adsorption of serum proteins that endogenously traffic to the lungs.^41–43^ Most of the injected mCherry was likely cleared from circulation after 18 hours, although small tissue fluorescence of cloaked mCherry that was delivered in MC3 LNPs still persisted in the liver (**Fig. 6c,d** and **Supplementary Fig. S8b**). As expected, systemic injection of free mCherry protein resulted in strong fluorescence only in the kidneys of mice (**Fig. 6b-d** and **Supplementary Fig. S8b**). As a moderately-sized protein (∼27 kDa), mCherry roughly falls within the glomerular filtration limit and thus is expected to be rapidly cleared from systemic circulation. Overall, the results from these experiments demonstrate that encapsulation of protein in LNPs decreases renal clearance and increases distribution to major organs, such as the liver and lungs, compared to that of free protein alone.

## Discussion

In this work, we present a facile bioconjugation strategy for protein delivery with cationic lipids that can be readily applied to virtually any protein cargo. By applying the anionic cloaking strategy on the selected proteins in this study, which vary widely in molecular weight (∼15 kDa to 150 kDa) and surface charge (pI less than 5 and greater than 8), we demonstrate the generalizability of this platform to enable highly efficient delivery into cells using clinically validated LNP formulations. The ability to exogenously introduce proteins into cells presents an immense opportunity to directly manipulate biological functions and has the potential to translate protein therapies that act on intracellular targets.

Initially, we demonstrate the efficacy of anionic cloaking using sfGFP as a model protein, illustrating that this approach enhances intracellular uptake efficiency when delivered with a commercial transfection reagent, and notably, even more so when combined with clinically validated LNPs. Our findings with sfGFP illustrate three key points of our delivery strategy: 1) anionic cloaking is necessary for protein delivery with commercial lipid reagents and LNPs, 2) successful protein internalization with LNPs requires the use of modified formulations supplemented with permanently cationic lipids, and 3) modification of protein with anionic moieties of different pK_a_ ‘s impacts LNP encapsulation and delivery efficiency. Our investigations involving RNase A and anti-β-catenin IgGs provide clear evidence of successful protein bioactivity following delivery, thereby validating our anionic cloaking strategy for the cytosolic delivery of diverse protein cargo following endosomal escape. Our studies also demonstrate the importance of redox-mediated disulfide cleavage and self-immolation of the cloaked sites for recovery of protein activity. In cases where protein function is impaired upon anionic cloaking, this provides an exciting oppportunity to control kinetics of cleavage (i.e., disulfide linkers that vary in electron-donating/withdrawing pendant groups) for sustained release of protein activity.^44^ Additionally, spatial control over protein activity could be achieved through organelle- and tissue-specific linker cleavage mechanisms (i.e., protease-specific cleavable linkers).^44^

Recent advances in protein delivery approahces have also investigated charge-based complexation of proteins with cationic lipids. The majority of these strategies, however, rely on genetically encoding anionic polypeptides into the backbones of protein cargos or using naturally anionic protein complexes to facilitate complexation with lipid reagents.^23–27^ The key advantage of our anionic cloaking strategy lies in the simplicity of its use – a broad-spectrum reagent that rapidly remodels the surface charge of any protein through the addition of sulfonate moeities, a chemical group that is not present in the toolkit of canonical amino acids.

Notably, in contrast to genetically encoded anionic polypeptides, our anionic tags facilitates a uniform, statistically distributed modification of accessible surface residues. The inherent self-immolative bond enables the modified protein to revert to its native form following cytosolic delivery, eliminating the need to navigate around critical protein sites for function. As previously alluded to, this feature not only alleviates constraints related to avoiding modification at crucial functional sites but also opens avenues for modulating protein activity based on the release kinetics of the anionic tags. Moreover, owing to the bioconjugation nature of this delivery approach, our strategy is versatile enough to be applied to functional proteins (such as antibodies) and proteins with post-translational modifications, all without requiring prior knowledge of their sequence or modification site. Looking ahead, we believe that our versatile delivery platform holds the potential to repurpose a wide range of commercial and therapeutic proteins for novel intracellular applications.

