## Supplementary file for "Bioreversible anionic cloaking enables intracellular protein delivery with ionizable lipid nanoparticles"

### MATERIALS

The following chemical reagents were used as received: Acryloyl chloride (Sigma Aldrich), triethylamine (Acros Organics), allylamine (Sigma Aldrich), diallylamine (Sigma Aldrich), propargylamine (Sigma Aldrich), *N*-Boc-ethylenediamine (Combi-Blocks), 1,8-Diazabicyclo[5.4.0]undec-7-ene (DBU; Sigma Aldrich), sodium 3-mercapto-1-propanesulfonate (3MPS; Sigma Aldrich), 4-mercaptobutyric acid (4MBA; A2B Chem), 2,2-Dimethoxy-2-phenylacetophenone (DMPA; Sigma-Aldrich), trifluoroacetic acid (TFA; BeanTown), 4-nitrophenyl chloroformate (Oakwood), bis(2-hydroxyethyl) disulfide (Sigma Aldrich), and D,L-dithiothreitol (DTT; Sigma Aldrich). DLin-MC3-DMA was purchased from MedChemExpress. SM-102 was purchased from Cayman Chemical. ALC-0315 was purchased from Echelon Biosciences. Cholesterol was purchased from Sigma Aldrich. 1,2-distearoyl-sn-glycero-3-phosphocholine (DSPC), 1,2-dioleoyl-3-trimethy-lammonium-propane (DOTAP), 1,2-dimyristoyl-rac-glycero-3-methoxypolyethylene glycol-2000 (PEG-DMG-2000) and distearoyl-rac-glycerol-PEG2K (PEG-DSG-2000) were purchased from Avanti Polar lipids. Lipofectamine 2000, Slide-A-Lyzer MINI Dialysis Devices (3.5 kDa MWCO), Hoechst 33342, and 5/6-carboxyfluorescein succinimidyl ester (NHS-Fluorescein) were purchased from Thermo Fisher Scientific. Micro Float-A-Lyzer Dialysis Devices (100 kDa MWCO) were purchased from Repligen. RNase A (Novagen, 10 mg/mL) was purchased from Sigma Aldrich. RNaseAlert Kit was purchased from Integrated DNA Technologies (IDT). CellTiter 96 Aqueous Non-Radioactive Cell-Proliferation Assay (MTS) and Dual-Glo Luciferase Assay were purchased from Promega. Monoclonal anti-β-catenin antibody (CAT-5H10), mouse IgG1 isotype control antibody (02-6100), and mouse anti-human IgG1, AlexaFluor 488 (A-10631) were purchased from Thermo Fisher Scientific.

### METHODS

#### Chemical synthesis of p-nitrophenyl carbonate compounds

For all synthesized materials, NMR spectroscopy was conducted on a Bruker 500 MHz NMR spectrometer. Liquid chromatography-mass spectrometry (LC-MS) was carried out on an Agilent 1200 Series LC/Thermo Fisher LTQ XL MSD equipped with an Agilent EC C18, 2.7 µm, 120 Å LC column (3 x 100 mm, reversed phase), UV diode-array detector monitoring 210 nm, 230 nm, 260 nm, 360 nm, and 505 nm wavelengths, and Agilent multimode source. Water with 0.1% formic acid (solvent A) and acetonitrile with 0.1% formic acid (solvent B) were used as LC-MS eluents. Compounds were eluted at a flow rate of 0.6 mL/min with a linear gradient of 5% to 100% solvent B over 10 minutes, then constant at 100% solvent B for 2 minutes before equilibrating the column back to 5% solvent B over 3 min. Masses were detected in either positive or negative ion mode. HPLC purification was performed on a 1100 Series Agilent HPLC system equipped with a UV diode array detector and a 1100 Infinity analytical scale fraction collector using reverse phase C18 column (4.6 x 150 mm, 5 *µ*m). The column compartment was kept at 25°C during fractionation. Solvents for HPLC were water with 0.1% trifluoroacetic acid (solvent A) and acetonitrile with 0.1% trifluoroacetic acid (solvent B). Compounds were eluted at a flow rate of 1 mL/min with 5% solvent B, followed by a linear gradient of 5% to 100% solvent B over 30 min, and finally 100% solvent B for 5 min before equilibrating the column back to 5% solvent B over 5 min.

*1. Synthesis of acrylamide monomers*

Triethylamine (TEA; 1.01 g, 11 mmol) and amines corresponding to each respective acrylamide monomer (refer below for specific amounts) were added to a flask with dichloromethane (DCM) (10 mL). Acryloyl chloride (0.91 g, 10 mmol) was added dropwise to the reaction mixture and stirred at 0°C for 15 min and then at room temperature for 1 h. The reaction was quenched with water, and the extracted organic layers were washed with 1 M HCl and brine. The organic layer was then dried over anhydrous $Na_2SO_4$. Excess DCM was dried in a rotary evaporator. The final product was purified by normal phase flash column chromatography with a gradient 20-60% ethyl acetate (EtOAc) in hexane and dried overnight in a vacuum concentrator.

*1.1. Synthesis of N-allylacrylamide*

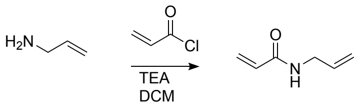

Amount of allylamine used was 0.57 g (10 mmol). Yield = 80%. $^1$H NMR (500 MHz, $CDCl_3$, δ): 6.30 (dd, *J* = 16.9, 1.4 Hz, 1H), 6.11 (dd, *J* = 17.0, 10.3 Hz, 1H), 5.93 – 5.79 (m, 1H), 5.66 (dd, *J* = 10.3, 1.4 Hz, 1H), 5.26 – 5.09 (m, 2H), 3.97 (dd, *J* = 11.7, 3.0 Hz, 2H).

*1.2. Synthesis of N,N-diallylacrylamide*

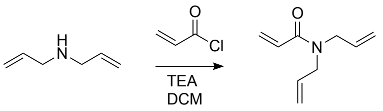

Amount of diallylamine used was 0.97 g (10 mmol). Yield = 75%. $^1$H NMR (500 MHz, $CDCl_3$, δ): 6.51 (dd, J = 16.8, 10.4 Hz, 1H), 6.39 (dd, J = 16.7, 2.2 Hz, 1H), 5.90 – 5.74 (m, 2H), 5.70 (dd, J = 10.4, 2.1 Hz, 1H), 5.27 – 5.13 (m, 4H), 4.01 (dd, J = 53.7, 5.3 Hz, 4H).

*1.3. Synthesis of N-propargylacrylamide*

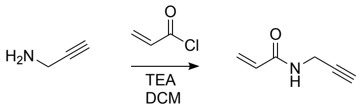

Amount of propargylamine used was 0.55 g (10 mmol). Yield = 78%. $^1$H NMR (500 MHz, $CDCl_3$, δ): 8.53 (s, 1H), 6.21 (dd, J = 17.2, 10.1 Hz, 1H), 6.11 (dd, J = 17.1, 2.1 Hz, 1H), 5.63 (dd, J = 10.1, 2.3 Hz, 1H), 3.94 (dd, J = 5.6, 2.5 Hz, 2H), 3.13 (t, J = 2.5 Hz, 1H).

*2. Synthesis of N-Boc-bis(propanamide)-ethylenediamine intermediates*

Acyrlamide monomers (refer below for specific amounts) were dissolved in MeOH (0.5 mL). *N*-Boc-ethylenediamine (160 mg, 1 mmol) and 1,8-Diazabicyclo[5.4.0]undec-7-ene (DBU; 152 mg, 1 mmol) were added to the solution, and the reaction mixture was stirred at 60°C for 48 h. The product was extracted with EtOAc, and the combined organic layers were washed with 1 M HCl and brine. The organic layer was then dried over anhydrous $Na_2SO_4$. Excess MeOH was dried in a rotary evaporator. The final product was purified by normal phase flash column chromatography with a gradient of 0-20% MeOH in DCM and dried overnight in a vacuum concentrator.

*2.1. Synthesis of N-Boc-bis(N-allylpropanamide)-ethylenediamine*

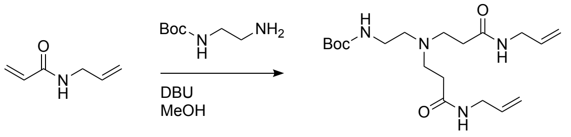

Amount of *N*-allylacrlyamide used was 244 mg (2.2 mmol). Yield = 42%. $^1$H NMR (500 MHz, $CDCl_3$, $\delta$): 6.53 (s, 2H), 5.89 – 5.77 (m, 2H), 5.16 (dd, *J* = 30.7, 13.8 Hz, 4H), 3.86 (t, *J* = 5.8 Hz, 4H), 3.21 – 3.13 (m, 2H), 2.71 (t, *J* = 5.8 Hz, 4H), 2.49 (t, *J* = 6.1 Hz, 2H), 2.33 (t, *J* = 6.0 Hz, 4H), 1.44 (s, 9H). LCMS (m/z): expected $[M]^+$: 382.26; observed $[M+H]^+$: 383.25.

*2.2. Synthesis of N-Boc-bis(N,N-diallylpropanamide)-ethylenediamine*

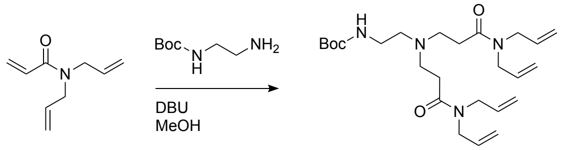

Amount of *N,N*-diallylacrylamide used was 332 mg (2.2 mmol). Yield = 35%. $^1$H NMR (500 MHz, $CDCl_3$, $\delta$): 5.84 – 5.70 (m, 4H), 5.26 – 5.09 (m, 8H), 4.13 (dd, *J* = 7.2, 7.2 Hz, 8H), 4.03 – 3.86 (m, 8H), 2.80 (t, *J* = 6.9 Hz, 2H), 2.44 (t, *J* = 6.9 Hz, 2H), 1.45 (s, 9H). LCMS (m/z): expected $[M]^+$: 462.32; observed $[M+H]^+$: 463.30.

*2.3. Synthesis of N-Boc-bis(N-(prop-2-ynyl)lpropanamide)-ethylenediamine*

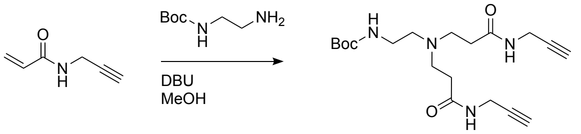

Amount of *N*-propargylacrylamide used was 240 mg (2.2 mmol). Yield = 50%. $^1$H NMR (500 MHz, $CDCl_3$, $\delta$): 6.85 (s, 2H), 4.07 (dd, *J* = 5.5, 2.6 Hz, 4H), 3.25 – 3.05 (m, 2H), 2.77 – 2.66 (m, 4H), 2.48 (t, *J* = 5.9 Hz, 2H), 2.37 – 2.31 (m, 4H), 2.25 (t, *J* = 2.6 Hz, 2H), 1.45 (s, 9H). LCMS (m/z): expected $[M]^+$: 378.23; observed $[M+H]^+$: 379.20.

<u>3. Synthesis of amine-terminated sulfonated and carboxylated intermediates</u>

Sodium 3-mercapto-1-propanesulfonate (3MPS) or 4-mercaptobutyric acid (4MBA) was dissolved in MeOH to prepare a 300 mM solution (refer below for specific amounts). 100 mg of *N*-Boc-bis(propanamide)-ethylenediamine and 2,2-Dimethoxy-2-phenylacetophenone (DMPA; 10%, wt/wt of total mass) were added to the solution. The reaction mixture was subjected to UV irradiation for 30 minutes at 20 mW/cm$^2$. The product was collected in cold ether, and the recovered precipitate was purified using RP-HPLC and lyophilized overnight. The purified intermediate was dissolved in 1 mL solution of 1:1 (v/v) DCM/trifluoroacetic acid (TFA) and stirred at room temperature for 1 h. The final product was obtained by drying in a rotary evaporator.

*3.1. Synthesis of amine-terminated SL2a*

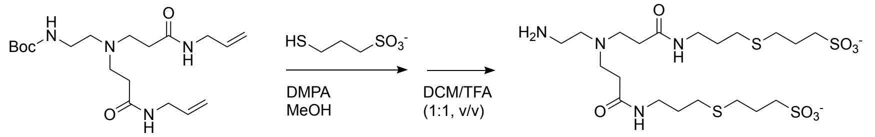

100 mg (0.26 mmol) of *N*-Boc-bis(*N*-allylpropanamide)-ethylenediamine and 186 mg (1.04 mmol, 4 eq.) of 3MPS dissolved in 3.49 mL of MeOH (300 mM) were used. Yield = 40%. $^1$H NMR (500 MHz, DMSO-$d_6$, δ): 3.26 – 3.22 (m, 4H), 2.89 – 2.77 (m, 4H), 2.57 – 2.53 (m, 14H), 2.48 – 2.43 (m, 4H), 1.85 – 1.78 (m, 10H). LCMS (m/z): expected $[M+H]^-$: 593.17; observed $[M+H]^-$: 593.18.

*3.2. Synthesis of amine-terminated SL2b*

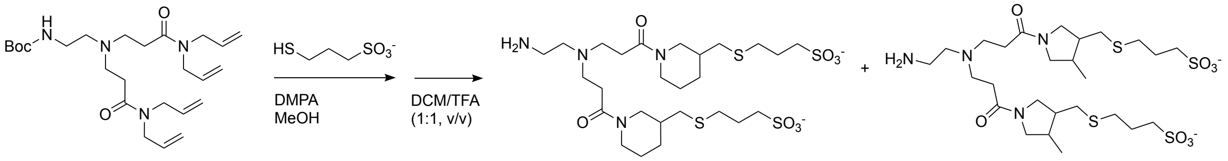

100 mg (0.22 mmol) of *N*-Boc-bis(*N,N*-diallylpropanamide)-ethylenediamine and 154 mg (0.85 mmol, 4 eq.) of 3MPS dissolved in 2.89 mL of MeOH (300 mM) were used. Yield = 43%. $^1$H NMR (500 MHz, DMSO-$d_6$, δ): 3.17 – 3.04 (m, 6H), 2.87 – 2.76 (m, 6H), 2.65 – 2.57 (m, 8H), 2.54 – 2.53 (m, 8H), 1.94 – 1.73 (m, 8H), 1.22 – 1.13 (m, 2H), 1.06 – 0.84 (m, 8H). LCMS (m/z): expected $[M+H]^-$: 673.24; observed $[M+H]^-$: 673.25.

*3.3. Synthesis of amine-terminated SL4*

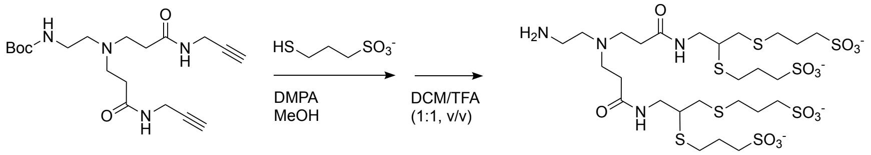

100 mg (0.26 mmol) of *N*-Boc-bis(*N*-(prop-2-ynyl)lpropanamide)-ethylenediamine and 377 mg (2.12 mmol, 8 eq.) dissolved in 7.05 mL of MeOH (300 mM) were used. Yield = 25%. $^1$H NMR (500 MHz, DMSO-$d_6$, δ): 3.54 – 3.17 (m, 10H), 2.92 – 2.84 (m, 2H), 2.79 – 2.69 (m, 4H), 2.68 – 2.57 (m, 12H), 2.55 – 2.53 (m, 12H), 1.87 – 1.75 (m, 6H). LCMS (m/z): expected $[M+3H]^-$: 901.11; observed $[M+2H]^{2-}$: 450.17; $[M+3H]^-$: 901.14.

*3.4. Synthesis of amine-terminated CL4*

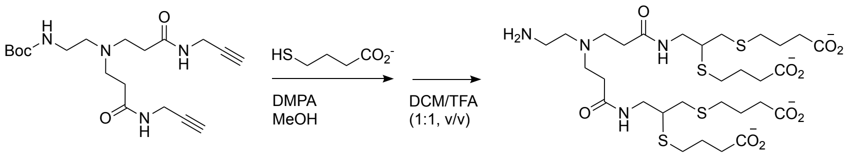

100 mg (0.26 mmol) of *N*-Boc-bis(*N*-(prop-2-ynyl)lpropanamide)-ethylenediamine and 245 mg of 4MBA (2.12 mmol, 8 eq.) and was dissolved in 7.05 mL of MeOH to prepare a 300 mM solution. Yield = 35%. $^1$H NMR (500 MHz, DMSO-$d_6$, δ): 2.97 – 2.85 (m, 4H), 2.74 – 2.69 (m, 4H), 2.67 – 2.60 (m, 6H), 2.57 – 2.52 (m, 10H), 2.40 – 2.19 (m, 12H), 1.93 – 1.80 (m, 4H), 1.78 – 1.69 (m, 6H). LCMS (m/z): expected $[M+3H]^-$: 757.24; observed $[M+3H]^-$: 757.26.

*4. Synthesis of sulfonated and carboxylated p-nitrophenyl carbonate compounds*

Amine-terminated sulfonated and carboxylated intermediates were dissolved in 1 mL of DMSO to prepare a 25 mM solution (refer below for specific amounts). Bis-*p*-nitrophenyl carbonate compounds and triethylamine (TEA) were added to the solution (refer below for specific amounts), and the reaction mixture was stirred at room temperature for 6 h. The final product was purified using RP-HPLC and lyophilized overnight.

*4.1. Synthesis of SL2a*

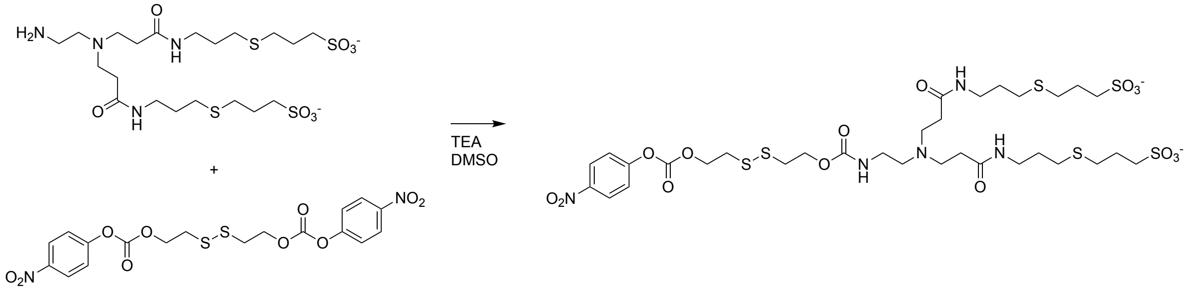

Amount of chemicals used were: 14.8 mg of amine-terminated SL2a (25 μmol), 10.9 mg of bis-*p*-nitrophenyl carbonate disulfide (22.5 μmol), and 5.05 mg of TEA (50 μmol). Yield = 47%. $^1$H NMR (500 MHz, DMSO-$d_6$, δ): 8.33 (d, *J* = 9.2 Hz, 2H), 7.58 (d, *J* = 9.2 Hz, 2H), 4.50 (t, *J* = 6.2 Hz, 2H), 4.23 (t, *J* = 6.4 Hz, 2H), 3.14 – 3.12 (m, 5H), 3.03 – 2.97 (m, 2H), 2.64 – 2.60 (m, 5H), 2.56 – 2.53 (m, 15H), 2.48 – 2.45 (m, 4H), 1.90 – 1.74 (m, 5H), 1.71 – 1.53 (m, 4H). LCMS (m/z): expected $[M+H]^-$: 938.19; observed $[M+H]^-$: 938.19.

*4.2. Synthesis of SL2b*

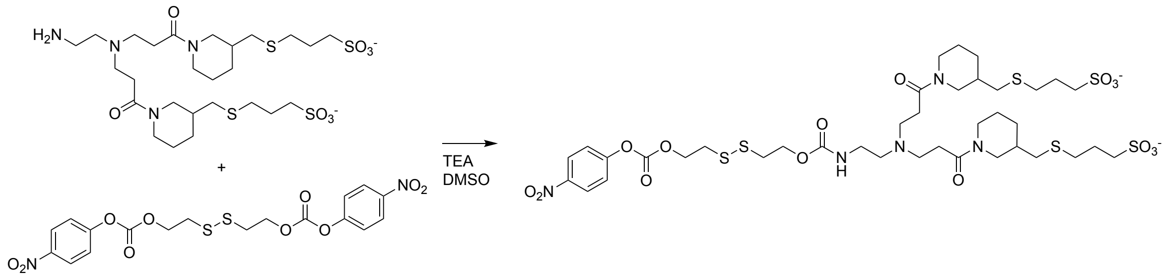

Although the amine-terminated SL2b is drawn as the 6-endo product, the starting material contained a mixture of the 6-endo and 5-exo isomers. Amount of chemicals used were: 16.8 mg of amine-terminated SL2b (25 μmol), 10.9 mg of bis-*p*-nitrophenyl carbonate disulfide (22.5 μmol), and 5.05 mg of TEA (50 μmol). Yield = 54%. $^1$H NMR (500 MHz, DMSO-$d_6$, δ): 8.33 (d, J = 9.2 Hz, 2H), 7.58 (d, J = 9.2 Hz, 2H), 4.50 (t, J = 6.2 Hz, 2H), 4.24 (t, J = 6.3 Hz, 2H), 3.12 (t, J = 6.2 Hz, 3H), 3.03 – 2.98 (m, 3H), 2.83 – 2.72 (m, 5H), 2.66 – 2.56 (m, 5H), 2.55 – 2.53 (m, 15H), 2.43 – 2.35 (m, 5H), 1.86 – 1.76 (m, 4H), 1.27 – 1.21 (m, 2H), 1.08 – 0.81 (m, 8H). LCMS (m/z): expected $[M+H]^{-}$: 1018.23; observed $[M+H]^{-}$: 1018.25.

#### *4.3. Synthesis of SL4*

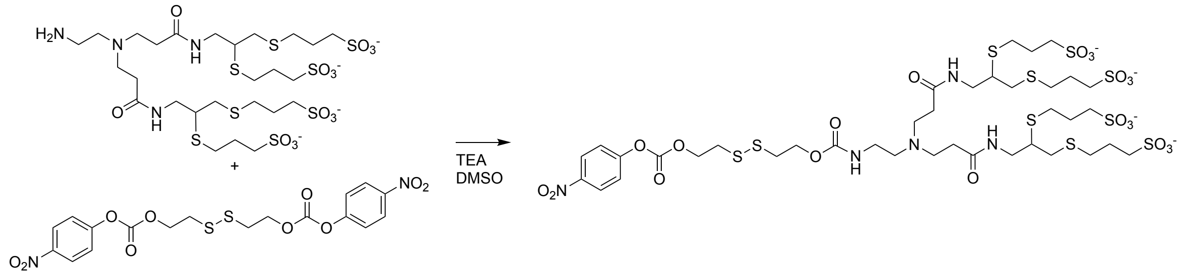

Amount of chemicals used were: 22.5 mg of amine-terminated SL4 (25 μmol), 10.9 mg of bis-*p*-nitrophenyl carbonate disulfide (22.5 μmol), and 5.05 mg of TEA (50 μmol). Yield = 40%. $^1$H NMR (500 MHz, DMSO-$d_6$, δ): 8.32 (d, J = 9.2 Hz, 2H), 7.59 (d, J = 9.2 Hz, 2H), 4.50 (t, J = 6.2 Hz, 2H), 4.24 (t, J = 6.5 Hz, 2H), 3.12 (t, J = 6.2 Hz, 2H), 3.00 (t, J = 6.5 Hz, 2H), 2.90 – 2.85 (m, 2H), 2.71 – 2.58 (m, 17H), 2.54 – 2.54 (m, 18H), 1.86 – 1.75 (m, 9H). LCMS (m/z): expected $[M+3H]^{-}$: 1246.11; observed $[M+H]^{3-}$: 414.71; $[M+2H]^{2-}$: 622.57; $[M+2Na+H]^{-}$: 1290.10.

#### *4.4. Synthesis of CL4*

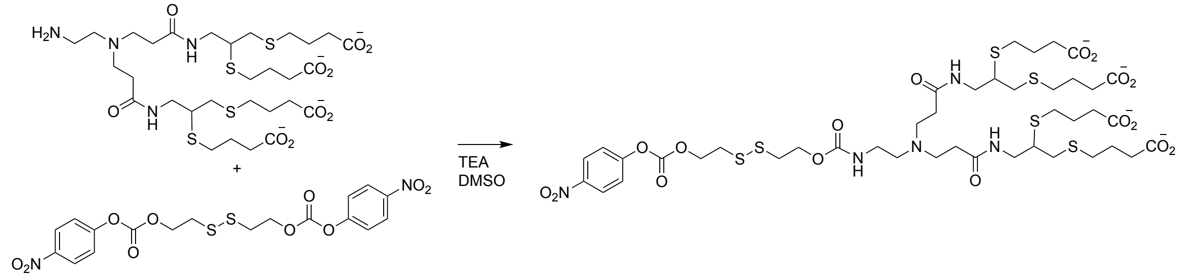

Amount of chemicals used were: 18.9 mg of amine-terminated CL4 (25 μmol), 10.9 mg of bis-*p*-nitrophenyl carbonate disulfide (22.5 μmol), and 5.05 mg of TEA (50 μmol). Yield = 35%. $^1$H NMR (500 MHz, DMSO-$d_6$, δ): 8.32 (d, J = 9.2 Hz, 2H), 7.59 (d, J = 9.8 Hz, 2H), 4.50 (t, J = 6.1 Hz, 2H), 4.24 (t, J = 6.7 Hz, 2H), 3.12 (t, J = 6.1 Hz, 2H), 3.00 (t, J = 5.8 Hz, 2H), 2.88 (t, J = 6.7 Hz, 2H), 2.75 – 2.57 (m, 18H), 2.54 – 2.53 (m, 18H), 1.85 – 1.76 (m, 8H). expected $[M+3H]^{-}$: 1102.24; observed $[M+3H]^{-}$: 1102.23.

**Protein expression and purification**

Plasmids encoding for sfGFP in pJL1 vector (Addgene 102634) and mCherry in pBAD vector (Addgene 54630) were transformed into competent *E.Coli* BL21(DE3) cells for protein production. For production of sfGFP, single colonies of BL21(DE3) cells containing pJL1-sfGFP were grown in 500 mL of Luria–Bertani (LB) medium containing kanamycin at 37°C. Expression was induced with 0.1 mM IPTG when $OD_{600}$ reached 0.6-0.8, and growth continued for 18 h at 16°C. For production of mCherry, single colonies of BL21(DE3) cells containing pBAD-mCherry were grown in 500 mL of LB medium containing ampicillin and supplemented with 0.2% D-glucose at 37°C for 8 hours. 500 mL of LB containing ampicillin were then added to the culture, and expression was induced with 0.2% L-arabinose for 18 h at 37°C. Cultures were harvested by centrifugation at 5000 *g* for 30 min at 4°C, and resuspended cell pellets were lysed with a high-pressure homogenizer (Avestin Emulsi-Flex C5). The insoluble fraction was cleared by centrifugation at 12000 *g* for 30 min at 4°C. 6xHis-tagged proteins were purified from clarified lysates by gravity flow using 750 µL HisPur Ni-NTA resin (Thermo Fisher) and washed with an endotoxin removal wash (Promega). Purified fractions were desalted into 1X PBS and concentrated (Pierce PES Protein Concentrators). Final protein concentration was determined by BCA assay (Thermo Fisher). Proteins were characterized by SDS/PAGE to confirm >95% purity.

**Protein bioconjugation and characterization**

For a typical procedure, *p*-nitrophenyl carbonate compounds were dissolved in DMSO to prepare a 10 mg/mL solution. Proteins were suspended in 10 mM HEPES buffer (pH 8.2) at a concentration of 1 mg/mL and incubated with *p*-nitrophenyl carbonate compounds overnight at 4°C. The mixtures were then transferred to a dialysis cup (Slide-A-Lyzer MINI Dialysis Devices, 3.5 kDA MWCO) and dialyzed against 1X PBS for 24 hours at 4°C to remove unreacted compounds. For experiments with fluorophore conjugated proteins, NHS-fluorescein was dissolved in DMSO to prepare a 10 mg/mL solution, and the same general reaction procedure was followed. First, proteins were labeled with NHS-fluorescein (2 molar eq.) overnight at 4°C, dialyzed for 24 hours at 4°C against 10 mM HEPES buffer (pH 8.2), and then reacted with *p*-nitrophenyl carbonate compounds at specified molar equivalencies. Final concentration of conjugated protein solutions was calculated from BCA assay (Thermo Fisher). Conjugated protein samples were characterized by MALDI-TOF-MS as well as by native gel electrophoresis (Bio-Rad) and isoelectric focusing (IEF; Thermo Fisher) as per manufacturers' protocols.

**Lipid nanoparticle (LNP) formulation and characterization**

Ionizable lipids (DLin-MC3-DMA, SM-102, and ALC-0315), DSPC, cholesterol, DMG-PEG-2000, DSG-PEG-2000, and DOTAP were dissolved in ethanol at defined molar ratios. Protein samples were suspended in 10 mM citrate buffer (at either pH 5 or pH 7.4) and pipette mixed rapidly into the ethanolic lipid solution at a volume ratio of 3:1 (proteins:lipids, v/v). The resulting mixture was dialyzed (Slide-A-Lyzer MINI Dialysis Devices, 3.5 kDA MWCO) against sterile 1X PBS for 2 h to remove ethanol. For *in vivo* biodistribution experiments, the formulation mixture was dialyzed (Micro Float-A-Lyzer, 100 kDa MWCO; Repligen) against sterile 1X PBS for 6 h. For experiments with sfGFP, RNase A, and antibodies, the molar ratio of ionizable lipid/DSPC/Chol/PEG-2000/DOTAP was 45/9/34.7/1.4/10 for all LNP formulations. For *in vivo* biodistribution experiments with mCherry, the molar ratio of MC3/DSPC/Chol/PEG-2000/DOTAP was 50/10/38.5/*X*/*Y*, where $X$ = 1.7 – 6.7 mol/mol (1.5 – 4.5 mol% PEG-2000) and $Y$ = 11.1 – 45 mol/mol (10 – 30 mol% DOTAP) for all MC3 LNP formulations. Particle size distribution and surface zeta potential of all LNP formulations were measured with a Malvern Zetasizer.

#### Encapsulation efficiency

LNP formulations of anionically-cloaked proteins were diluted with 1X PBS to a concentration of 50 μg/mL. 10 μL LNP samples (corresponding to 0.5 μg of total protein) was incubated in the presence or absence of 1% Triton-X (vol/vol), and Triton-X treated samples were sonicated for 5 minutes to lyse LNPs. Samples were mixed with 1 volume equivalent of native gel loading buffer and loaded onto 10% polyacrylamide gels (Bio-Rad) for native gel electrophoresis. Quantification of encapsulation efficiency was calculated from in-gel fluorescence intensities using the following formula: $(LNP_{lysed} - LNP_{intact})/LNP_{lysed} \times 100$. Image analysis and quantification was performed with Bio-Rad Image Lab software.

#### Cell lines and cell culture

HEK293T, HeLa, A549, SK-BR-3, SK-OV-3, and DLD-1 cells were procured from lab stocks. HEK293T, HeLa, and A549 cells were maintained in DMEM media containing 10% FBS and 1% penicillin/streptomycin. SK-BR-3 and SK-OV-3 cells were maintained in McCoy 5A media containing 10% FBS and 1% penicillin/streptomycin. DLD-1 cells were maintained in RPMI 1600 media containing 10% FBS and 1% penicillin/streptomycin. All cells were maintained at 37°C, 5% $CO_2$ and 90% relative humidity.

#### Cell viability assay

A total of 10,000 cells per well were plated in a clear 96-well tissue plate and incubated overnight at 37°C for delivery experiments. After experiment, media was aspirated, and cells were washed once with 1X PBS. 100 μL of FluoroBrite DMEM and 10 μL of MTS solution (Promega) were added to each well, and the plate was incubated for 1 hour. Absorbance measurements were taken at 490 nm on a TECAN Infinite M1000 Pro microplate reader and normalized to untreated cells (100%) or clear media (0%).

#### TOPFlash assay

A total of 10,000 DLD-1 cells per well were plated in a white-bottom 96-well tissue culture plate and incubated overnight at 37°C. The following day, 50 ng of either TOPFlash plasmid (Addgene 12456) or FOPFlash plasmid (TOPFlash mutant; Addgene 12457) and 5 ng of Renilla luciferase plasmid (Addgene 87121) was co-transfected into cells with Lipofectamine 2000 (Thermo Fisher) in OptiMEM reduced serum media as per manufacturer's protocol (0.2 μL Lipofectamine 2000/well). 6 hours after transfection, the transfection mixture was replaced with fresh RPMI 1600 media containing 10% FBS and 1% penicillin/streptomycin. MC3 LNPs formulated with antibodies were then added to each well and incubated for 24 hours 37°C. Following incubation, media was aspirated, and cells were washed once with 1X PBS. Cells were lysed, and the firefly and Renilla luminescence signals were measured sequentially by the Dual-Glo Luciferase Assay System (Promega) as per manufacturer's protocol. Plates were read on a TECAN Infinite M1000 Pro microplate reader. The luciferase activities were measured and normalized against the control Renilla activities.

#### *In vitro* protein delivery

For a typical protein delivery procedure, 75,000 cells were seeded in a 24-well tissue culture plate and incubated overnight at 37°C. For transfections with Lipofectamine 2000 (Thermo Fisher), 500 nM of protein diluted in 10 µL of OptiMEM reduced serum media, and 2 µL of Lipofectamine 2000 was suspended in 8 µL of OptiMEM. The protein and lipid reagent solutions were pipette mixed and incubated at room temperature for 10 minutes to allow for complexation. The resulting 20 µL transfection mixture was then added to cells containing 380 µL of cell culture media to a final volume of 400 µL. For transfections with LNPs, LNP solutions formulated with proteins were diluted in 75 µL of 1X PBS and added to 325 µL cell culture media to a final volume of 400 µL. Delivery experiments were performed at 37°C for 6 h before flow cytometric analysis (see below).

#### Flow cytometry

Following delivery experiments in a 24-well tissue culture plate, media was aspirated, and cells were washed three times with PBS. Cells were detached and harvested in 1X PBS by pipetting each well 10-15 times. Cells were pelleted by centrifugation at 500 *g* for 5 minutes, and the supernatant was aspirated and replaced with fresh 1X PBS for flow cytometric analysis. Flow cytometry was performed on a FACSCalibur (Bectron Dickinson) or Attune NxT (Thermo Fisher) flow cytometer. FlowJo Version 10 was used to analyze samples by geometric mean fluorescence determined from 10,000 events.

#### Confocal microscopy

Cells were plated at 10,000 cells/cm$^2$ on an 8-well chamber slide pretreated with bovine gelatin and incubated overnight at 37°C. Lipofectamine/sfGFP complexes or LNPs formulated with sfGFP were added to each chamber in a total volume of 200 µL of cell culture media. Delivery experiments were performed at 37°C for 6 h. Media was aspirated, and cells were washed three times with 1X PBS. 200 µL of FluoroBrite DMEM media containing Hoescht diluted 1:10,000 was added to each chamber. After 15 minutes of incubation, media was aspirated, and cells were washed three times with 1X PBS. Chambers were then refilled with 200 µL of FluoroBrite DMEM before imaging. Samples were imaged on an inverted Zeiss LSM88-confocal microscope (i880) using a 40×water immersion objective. Images were analyzed with FIJI software.

#### RNase A activity assay

Ribonuclease activity of RNase A and modified RNase A was assessed with RNaseAlert Kit (IDT). Briefly, 0.1 µg of RNase A samples diluted in 80 µL of nuclease free water was added to a black 96-well plate. To this solution, 10 µL RNaseAlert Substrate mixed with 10 µL of 10X assay buffer was added to each well. Fluorescence emission at 520 nm (490 nm excitation) was read on a TECAN Infinite M1000 Pro microplate reader using a kinetic cycle up to either 90 minutes or 6 hours.

**ELISA**

A 96-well enzyme immunoassay plate was coated with 100 µL of recombinant β-catenin protein (Sino Biological 11279-H07E) at 10 µg/mL in 0.05 M $NaCO_3$ buffer (pH 9.6) overnight at 4°C. The plate was washed three times with 200 µL of PBST (1X PBS and 0.1% Tween) per well and blocked with 200 µL of 1X PBS containing 3% milk per well overnight at 4°C. The plate was washed three times with 200 µL of PBST per well. Serial dilutions of anti-β-catenin antibody samples in blocking buffer were added at 50 µL per well, and the plate was slowly mixed for 1 h at room temperature. The plate was washed three times with 200 µL of PBST per well and then incubated with anti-mouse antibody conjugated to horsereadish peroxidase (HRP) diluted 1:10,000 in 50 µL PBST+1% milk for 1 h with slow mixing. The plate was washed three times with 200 µL of PBST per well before the addition of 50 µL of 1-Step Ultra TMB (Thermo Fisher). The reaction was allowed to incubate with slow mixing and then quenched with 50 µL of 3N $H_2SO_4$. Absorbance measurements were taken at 450 nm on a TECAN Infinite M1000 Pro microplate reader.

***In-vivo* biodistribution**

All animal experiments were approved by the Institution of Animal Care and Use Committees of Cornell University and were consistent with local, state, and federal regulations as applicable. Female SKH-1 mice (elite strain 477, 8-10 weeks old) were obtained from Charles River Laboratories. All mice were injected intravenously (IV) through the tail-vein. Before injections, total protein amount of MC3 LNPs formulated with mCherry was calculated from BCA assay of LNP samples. Mice weighing 18-20 g were IV injected with 1X PBS as well as free mCherry and MC3 LNPs formulated with mCherry at a dose of 1 mg/kg of total protein. Mice were euthanized, and *ex vivo* imaging of harvested organs was performed 1.5 h and 18 h post injection on an IVIS Spectrum system (Perkin Elmer). Images were analyzed with Living Image (Perkin Elmer) software.

**Statistical analysis**

Statistical analyses were performed by one-way ANOVA with Bonferroni correction for multiple comparisons, unpaired t-tests followed by Bonferroni-Dunn correction for multiple comparisons, or two-way ANOVA followed by Bonferroni correction for multiple comparisons as appropriate. Statistical analysis was performed on GraphPad Prism software, v10 (GraphPad). A P-value $<0.05$ was considered statistically significant.

SUPPLEMENTARY FIGURES

| | Z-average (nm) | PDI | Zeta potential (mV) |
|---|---|---|---|
| **2 wt/wt MC3 (pH 5)** | 273.7 ± 4.2 | 0.114 ± 0.015 | -0.48 ± 0.26 |
| **5 wt/wt MC3 (pH 5)** | 273.8 ± 7.4 | 0.132 ± 0.013 | -1.04 ± 1.16 |
| **10 wt/wt MC3 (pH 5)** | 247.7 ± 3.3 | 0.135 ± 0.019 | -1.43 ± 0.49 |
| **2 wt/wt MC3 +10 mol% DOTAP (pH 5)** | 290.6 ± 3.0 | 0.086 ± 0.014 | -0.45 ± 0.96 |
| **5 wt/wt MC3+10 mol% DOTAP (pH 5)** | 354 ± 6.9 | 0.175 ± 0.012 | -3.52 ± 1.16 |
| **10 wt/wt MC3+10 mol% DOTAP (pH 5)** | 294.6 ± 7.1 | 0.110 ± 0.024 | -2.56 ± 0.42 |
| **2 wt/wt MC3 (pH 7.4)** | 141.7 ± 4.0 | 0.140 ± 0.013 | -3.96 ± 0.28 |
| **5 wt/wt MC3 (pH 7.4)** | 181 ± 2.8 | 0.191 ± 0.020 | -3.43 ± 0.93 |
| **10 wt/wt MC3 (pH 7.4)** | 182.5 ± 3.2 | 0.187 ± 0.015 | -2.13 ± 0.29 |
| **2 wt/wt MC3 +10 mol% DOTAP (pH 7.4)** | 242.9 ± 3.2 | 0.129 ± 0.027 | 0.42 ± 0.81 |
| **5 wt/wt MC3+10 mol% DOTAP (pH 7.4)** | 239.2 ± 4.7 | 0.082 ± 0.014 | -1.28 ± 0.23 |
| **10 wt/wt MC3+10 mol% DOTAP (pH 7.4)** | 211.3 ± 5.5 | 0.121 ± 0.016 | -5.41 ± 1.46 |

**Supplementary Table S1. Dynamic light scattering of LNPs formulated with anionically-cloaked sfGFP.** Size (z-avg), PDI, surface zeta potential of MC3 LNPs formulated with sfGFP-SL4 (modified with 30 molar eq.). LNPs were formulated in pH 5 and pH 7.4 buffers and supplemented with or without 10 mol% DOTAP. All data are mean ± SD (*n* = 3).

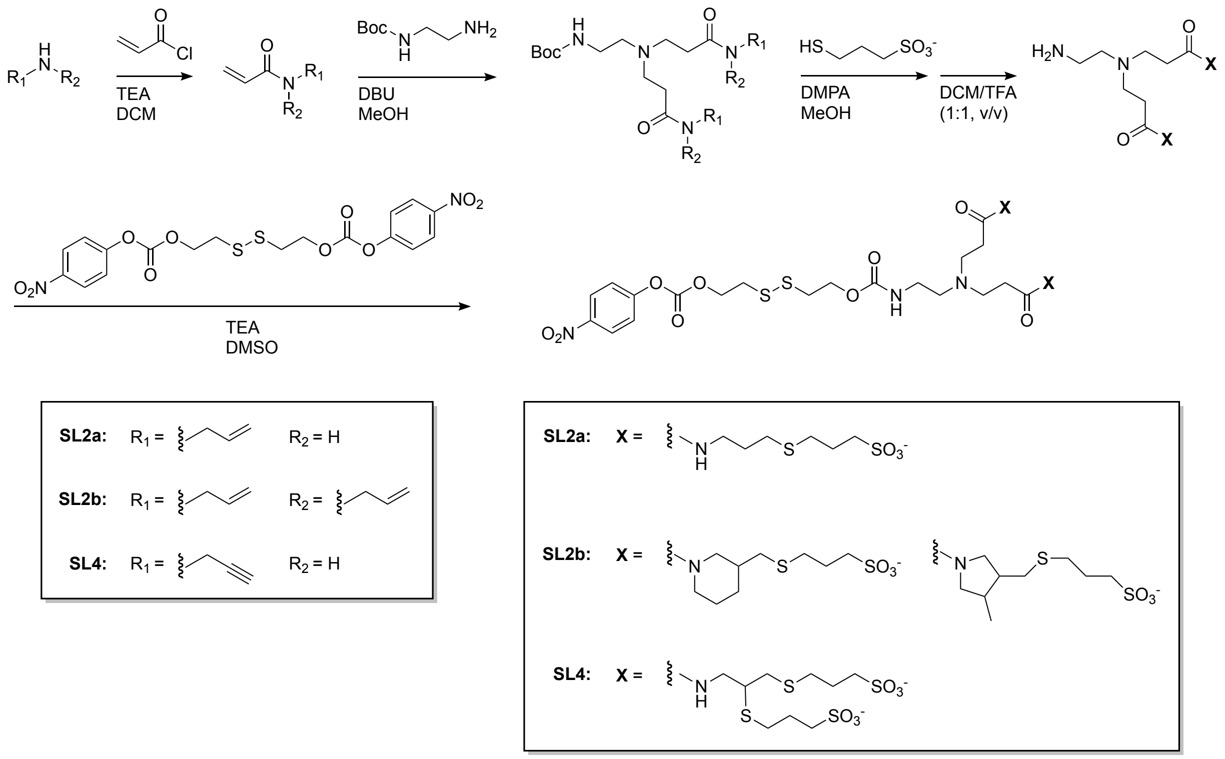

**Supplementary Figure S1. General scheme for the synthesis of anionic *p*-nitrophenyl carbonate compounds.** Scheme outlining general synthetic steps for producing SL2a, SL2b, and SL4.

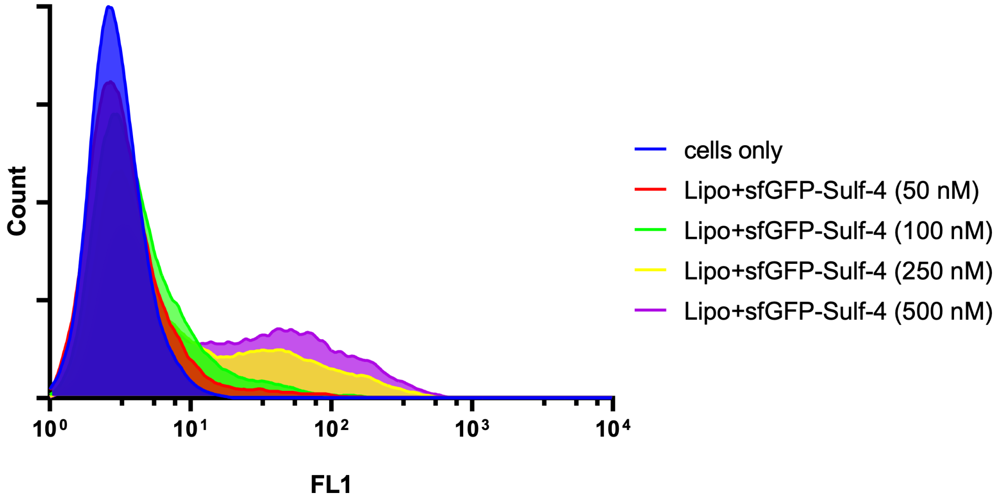

**Supplementary Figure S2. Intracellular delivery of anionically-cloaked sfGFP with Lipofectamine 2000.** Representative flow cytometry histograms of sfGFP (modified with 30 molar eq. of SL4) transfections into HEK293T cells with Lipofectamine 2000 from 50–500 nM.

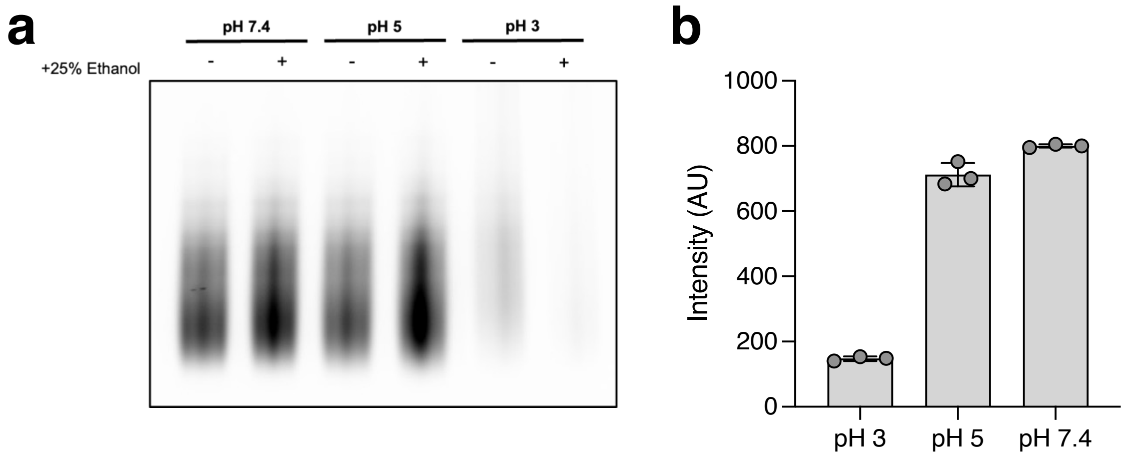

**Supplementary Figure S3. Effect of formulation pH on sfGFP activity.** (a) In-gel fluorescence images of gels run under native conditions of sfGFP-SL4 (modified with 30 molar eq.) incubated in different pH buffers and in the presence of ethanol (1 µg protein/well). Indicated conditions are intended to reflect the conditions of LNP formulation. (b) Emission spectrum of sfGFP-SL4 at varying pH conditions. All data are mean ± SD (*n* = 3).

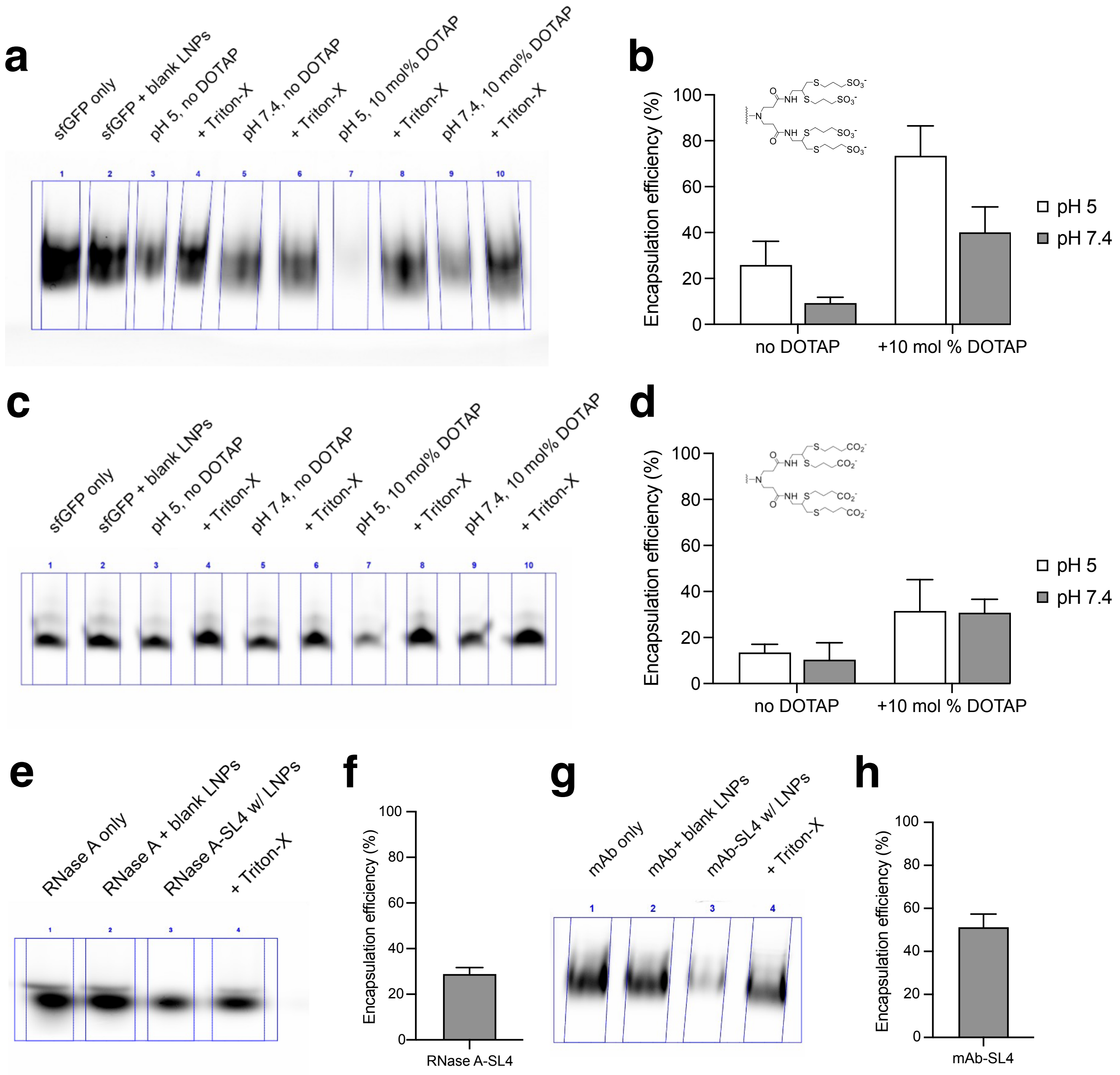

**Supplementary Figure S4. Encapsulation efficiency of anionically-cloaked proteins formulated with LNPs.** LNPs were formulated in pH 5 and pH 7.4 buffers and supplemented with or without 10 mol% DOTAP. LNP samples were treated with Triton-X to dissolve LNPs and release encapsulated proteins. Gels were loaded with 0.5 µg protein/well. Percent encapsulation was calculated by normalizing signal from LNP samples with signal from LNP samples treated with Triton-X. (a) Representative in-gel fluorescence image of gel electrophoresis run under native conditions of sfGFP-SL4 (modified with 30 molar eq. of SL4) formulated with MC3 LNPs (10 wt/wt, MC3/sfGFP). (b) Encapsulation efficiency of sfGFP-SL4 in MC3 LNPs as calculated from gel densitometry. (c) Representative in-gel fluorescence image of gel electrophoresis run under native conditions of sfGFP-CL4 (modified with 30 molar eq. of SL4) formulated with MC3 LNPs (10 wt/wt, MC3/sfGFP). (d) Encapsulation efficiency of sfGFP-SL4 in MC3 LNPs as calculated from gel densitometry. (e) Representative in-gel fluorescence image of gel electrophoresis run under native conditions of RNase A-SL4 (modified with 2 molar eq. of NHS-Fluorescein and 10 molar eq. of SL4) formulated with MC3 LNPs (10 wt/wt, MC3/RNase A) supplemented with 10 mol% DOTAP in pH 5 buffer. (f) Encapsulation efficiency of RNase A-SL4 in MC3 LNPs as calculated from gel densitometry. (g) Representative in-gel fluorescence image of gel electrophoresis run under native conditions of AlexaFluor488 secondary IgG-SL4 (modified with 30 molar eq. of SL4) formulated with MC3 LNPs (2 wt/wt, MC3/IgG) supplemented with 10 mol% DOTAP in pH 5 buffer. (h) Encapsulation efficiency of AlexaFluor488 secondary IgG-SL4 in MC3 LNPs as calculated from gel densitometry. All data are mean ± SD ($n = 3$).

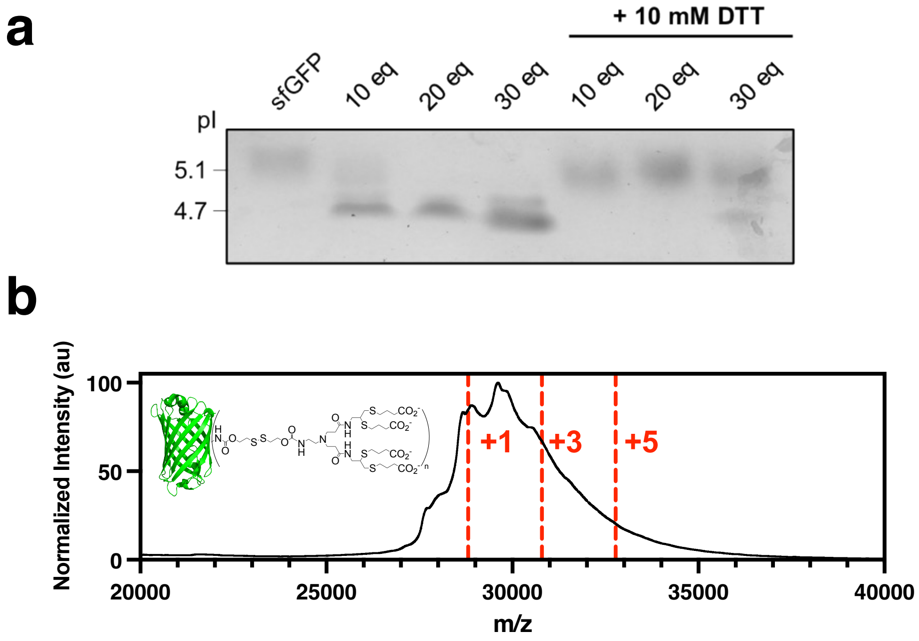

**Supplementary Figure S5. Conjugation and characterization of carboxylated sfGFP.** (a) IEF gel of sfGFP conjugated to varying molar eq. of CL4 before and after incubation with 10 mM DTT. (b) MALDI spectra of sfGFP conjugated to CL4 (modified with 30 molar eq.).

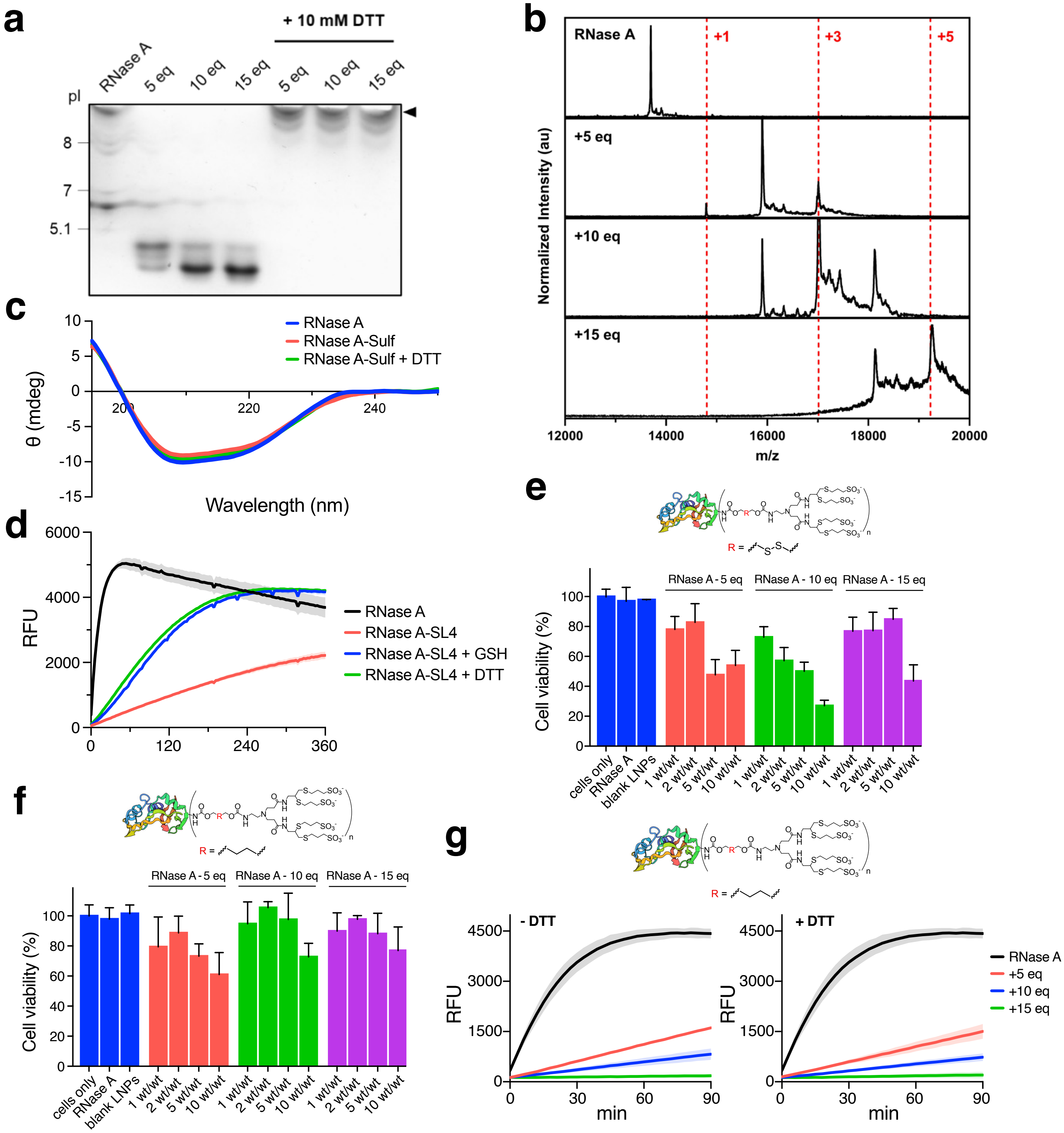

**Supplementary Figure S6. Characterization of anionically-cloaked RNase A.** (a) IEF gel of RNase A conjugated to varying molar eq. of SL4 before and after incubation with 10 mM DTT. (b) MALDI-MS spectra of RNase A and RNase A modified with varying molar eq. of SL4. (c) CD spectra of native RNase A, RNase A-SL4 (modified with 10 molar eq. of SL4), and RNase-SL4 incubation with 10 mM DTT. (d) RNase activity assay of native RNase A and RNase A modified with SL4 ($n = 4$). RNase-SL4 samples were co-incubated with 10 mM of either GSH or DTT during the assay. (e) Viability of HEK293T cells transfected with 500 nM of RNase A-SL4 as measured by MTS assay ($n = 4$). RNase A was modified with SL4 containing redox-cleavable disulfide linker and formulated with MC3 LNPs (1-10 wt/wt, MC3/RNase A) with 10 mol % DOTAP in pH 5 citrate buffer. RNase A transfections were performed for 48 hours. (f) Viability of HEK293T cells transfected with 500 nM of RNase A modified with non-cleavable SL4 as measured by MTS assay. RNase A was modified with non-cleavable SL4 containing butyl linker and formulated with MC3 LNPs (1-10 wt/wt, MC3/RNase A) with 10 mol % DOTAP in pH 5 citrate buffer. RNase A transfections were performed for 48 hours. (g) RNase activity assay of native RNase A and RNase A modified with non-cleavable SL4. Activity assays were repeated for RNase A samples incubated overnight with 10 mM DTT. All data are mean ± SD ($n = 4$ for MTS; $n = 4$ for ribonuclease assay).

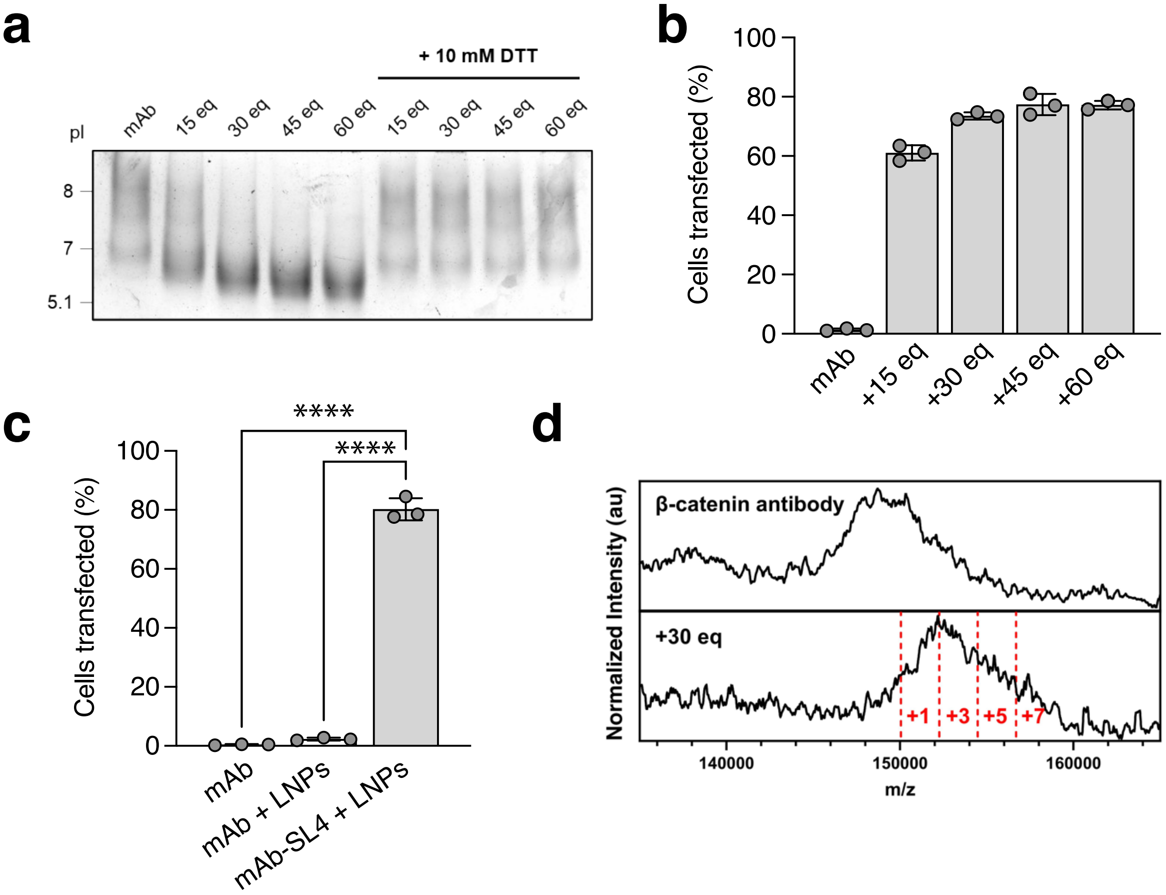

**Supplementary Figure S7. Preparation and delivery of anionically-cloaked IgG antibody.** Data shown are for MC3 LNPs (MC3/IgG, 2 wt/wt) supplemented with 10 mol% DOTAP and formulated at pH 5. (a) IEF gel of AlexaFluor 488 secondary IgG conjugated to varying molar eq. of SL4 before and after incubation with 10 mM DTT. (b) Percent IgG positive HEK293T cells following transfections of AlexaFluor 488 secondary IgG conjugated to varying molar eq. of SL4 and formulated with MC3 LNPs, as analyzed by flow cytometry. Transfections were performed for 6 hours. (c) Percent of AlexaFluor 488 IgG positive cells following 250 nM transfections into HEK293T cells. AlexaFluor 488 IgG-SL4 was cloaked with 30 molar eq. of SL4. Transfections were performed for 6 hours. Statistical significance was determined by unpaired t tests (*$p < 0.05$, **$p < 0.01$, ***$p < 0.001$, ****$p < 0.0001$). (d) MALDI spectra of anti-β-catenin IgG and IgG modified with 30 molar eq. of SL4. All data are mean ± SD ($n = 3$ for flow cytometry).

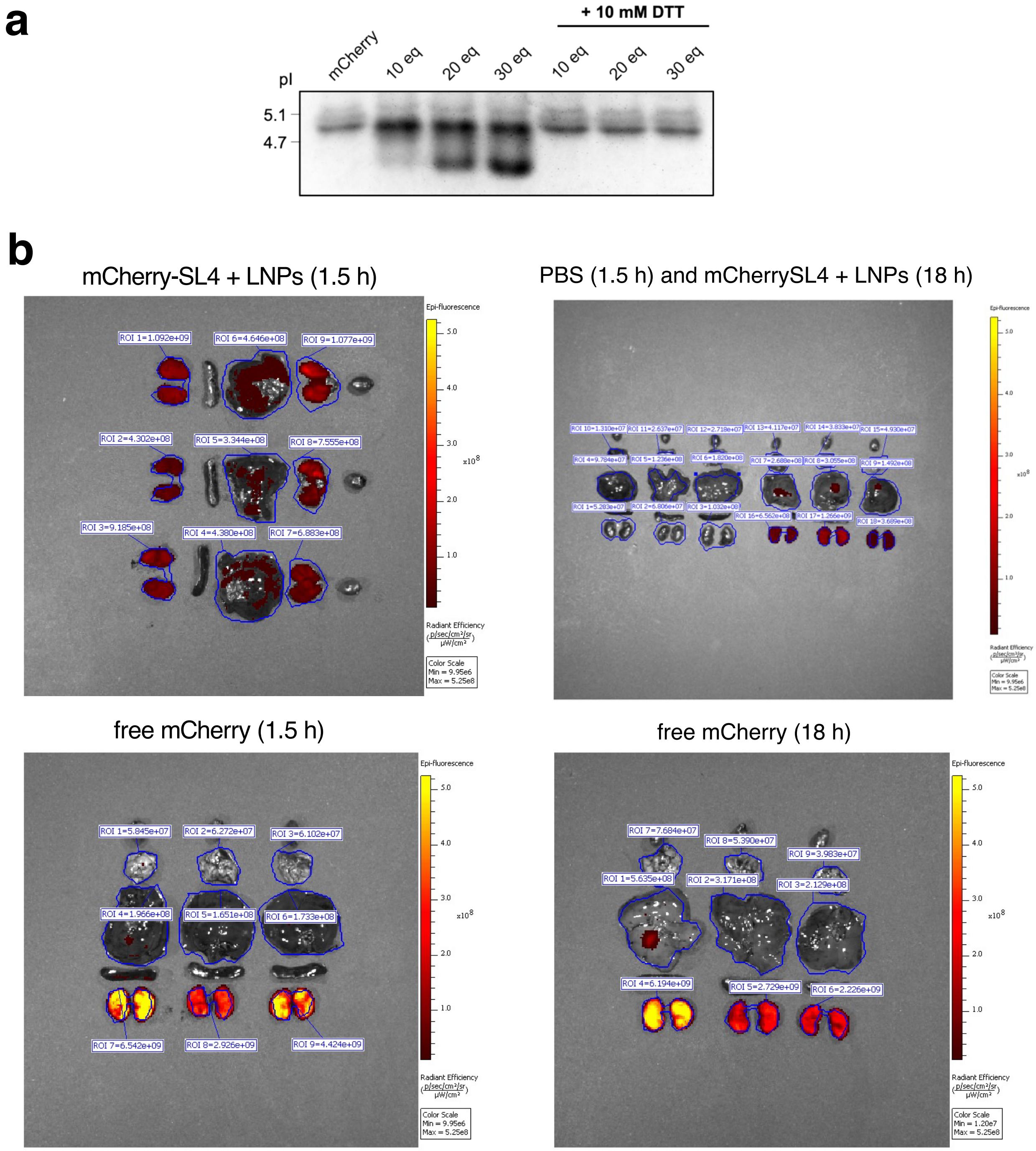

**Supplementary Figure S8. In vivo biodistribution of anionically-cloaked mCherry delivered with MC3 LNPs.** (a) IEF gel of mCherry conjugated to varying molar eq. of SL4 before and after incubation with 10 mM DTT. (b) *Ex vivo* fluorescent images of harvested organs following tail vein injections of SKH1 mice with PBS, free mCherry, and mCherry-SL4 formulated in MC3 LNPs. MC3 LNPs were formulated with 3 mol% PEG-DMG-2000 and 30 mol% DOTAP. Doses were 1 mg/kg of total protein. Images were taken 1.5 h and 18 h post-injection. Calculated radiant efficiencies of ROIs corresponding to lungs, liver, and kidneys are shown.

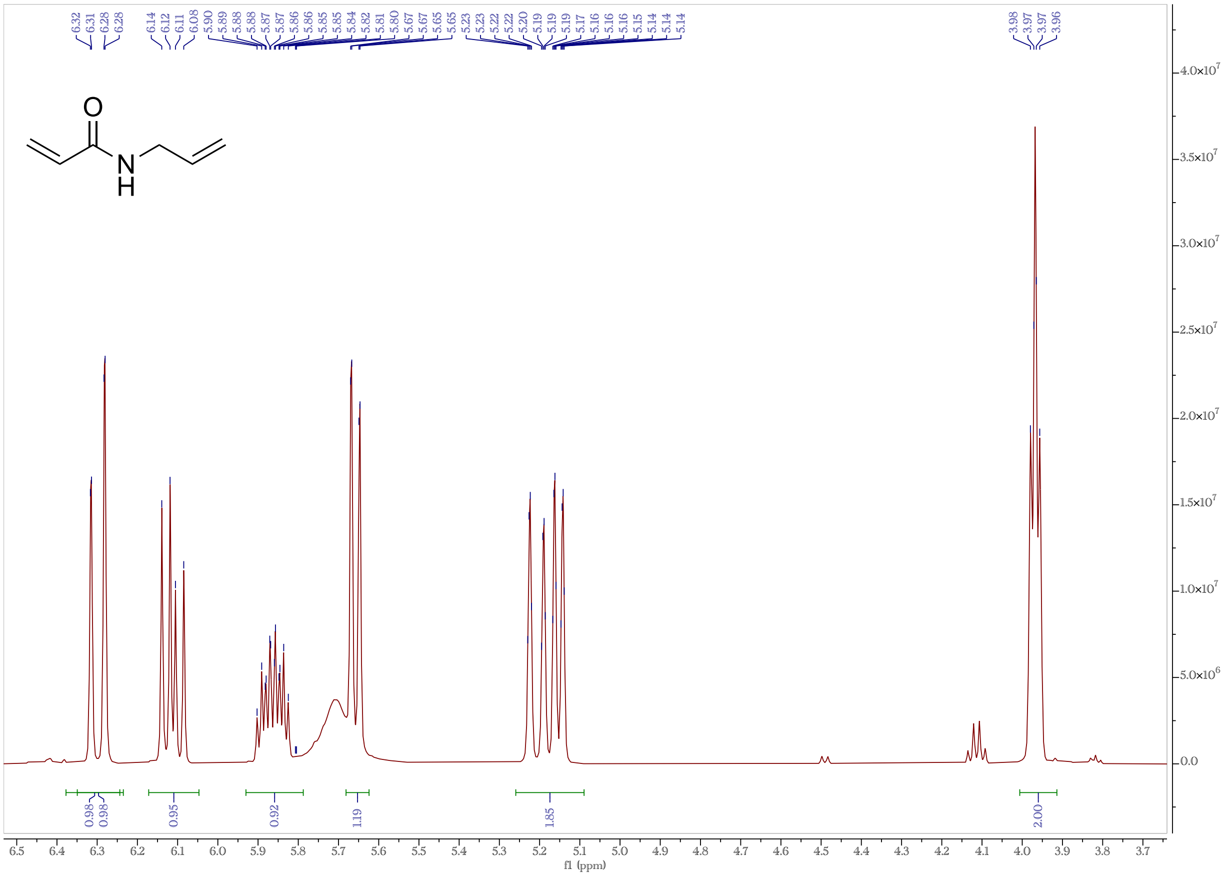

**Supplementary Figure S9.** $^1$H NMR (500 MHz, $CDCl_3$) of *N*-allylacrylamide.

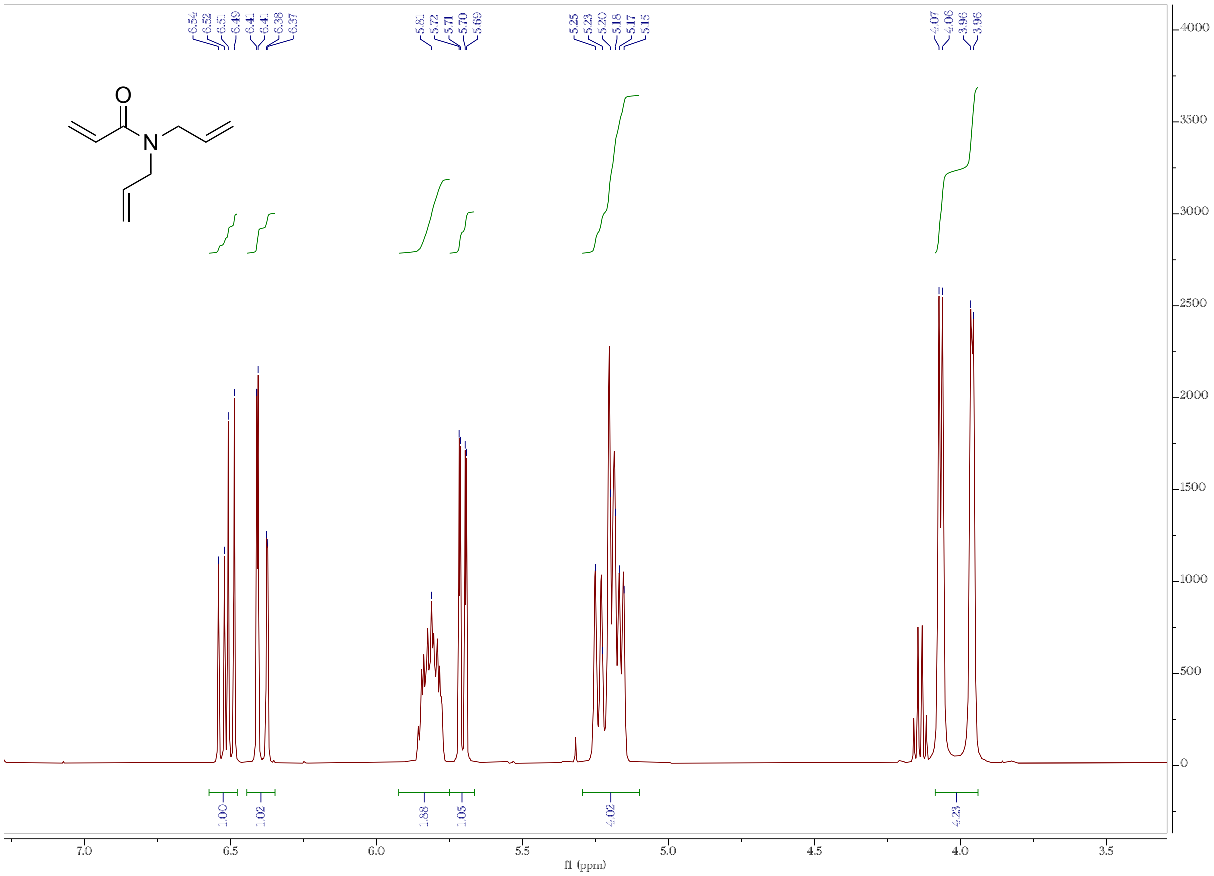

**Supplementary Figure S10.** $^1$H NMR (500 MHz, $CDCl_3$) of allylacrylamide.

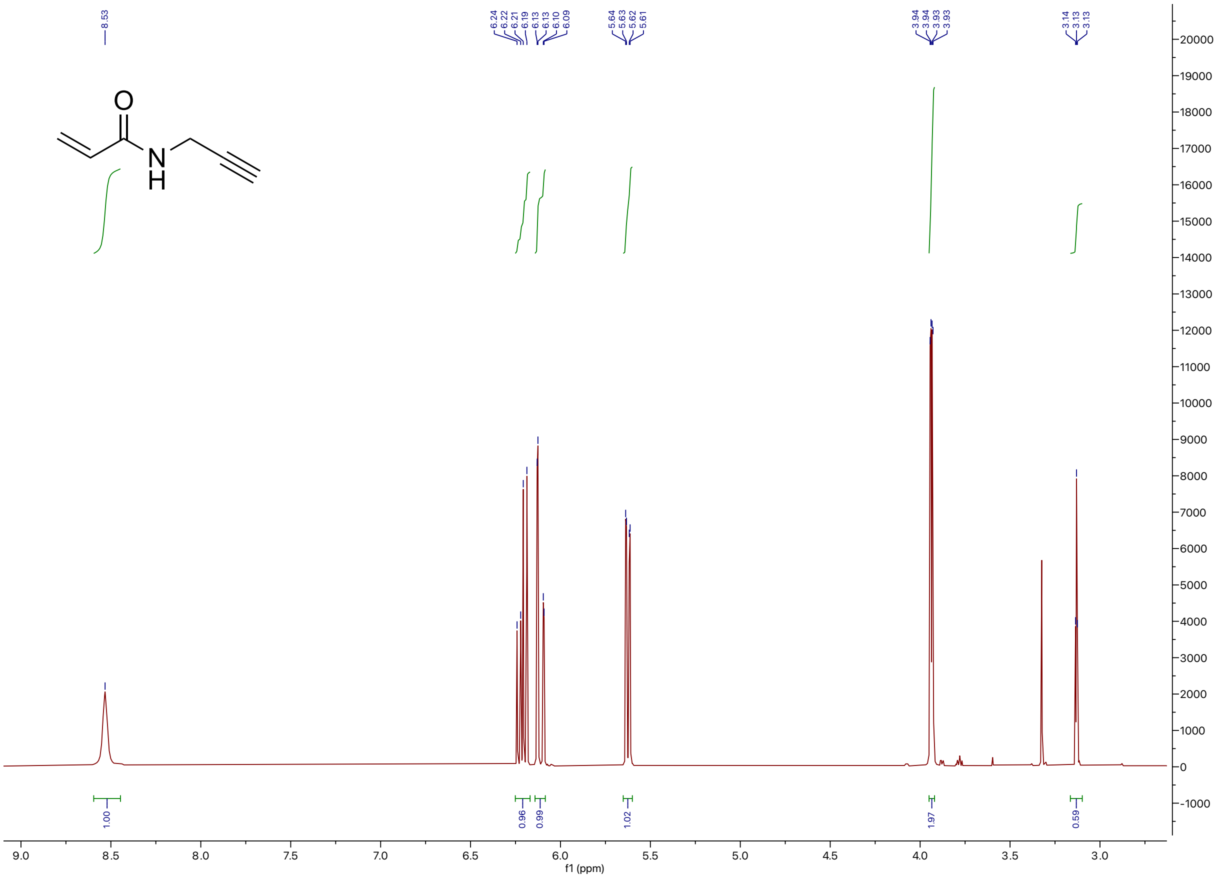

**Supplementary Figure S11.** $^1$H NMR (500 MHz, $CDCl_3$) of propargylacrylamide.

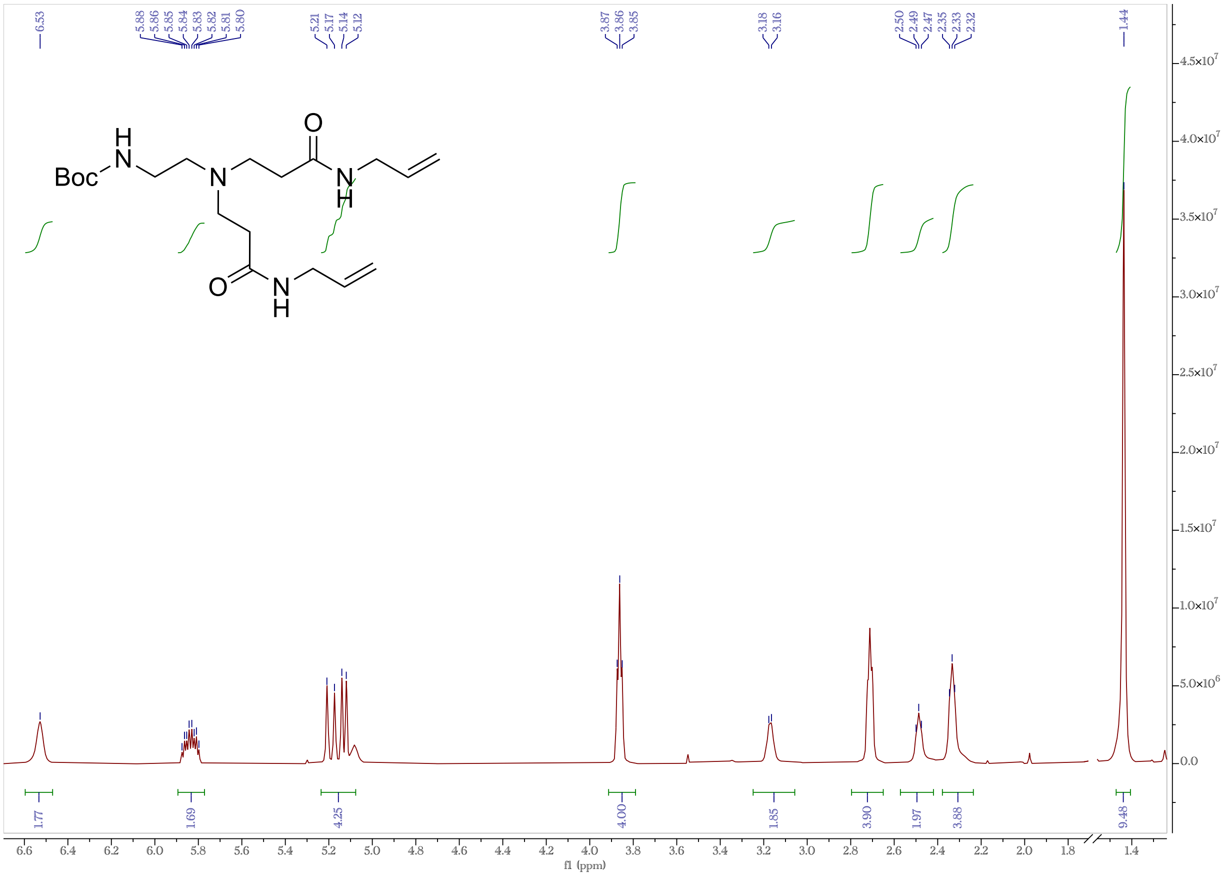

**Supplementary Figure S12.** $^{1}H$ NMR (500 MHz, $CDCl_3$) of *N*-Boc-bis(*N*-allylpropanamide)-ethylenediamine.

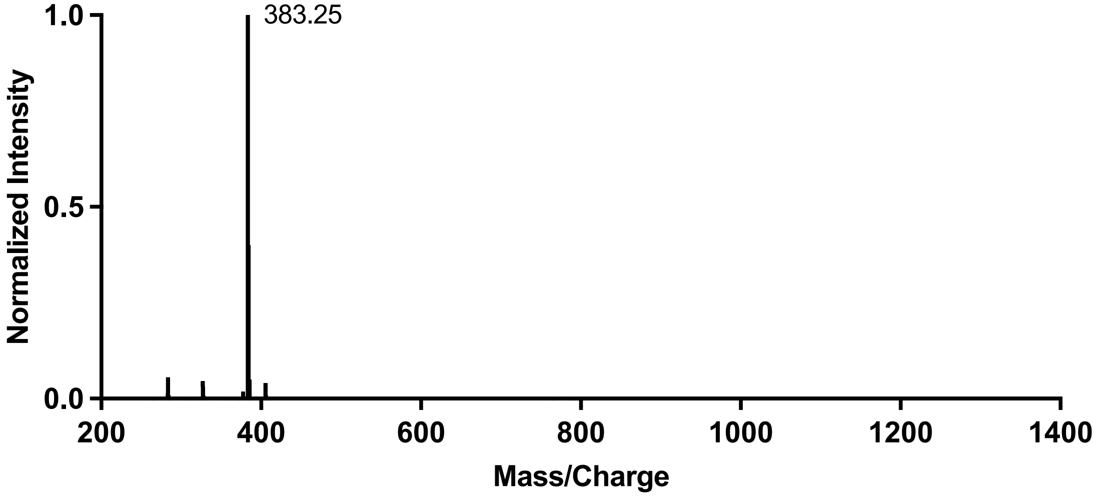

**Supplementary Figure S13.** LC/MS analysis of *N*-Boc-bis(*N*-allylpropanamide)-ethylenediamine (positive ion mode).

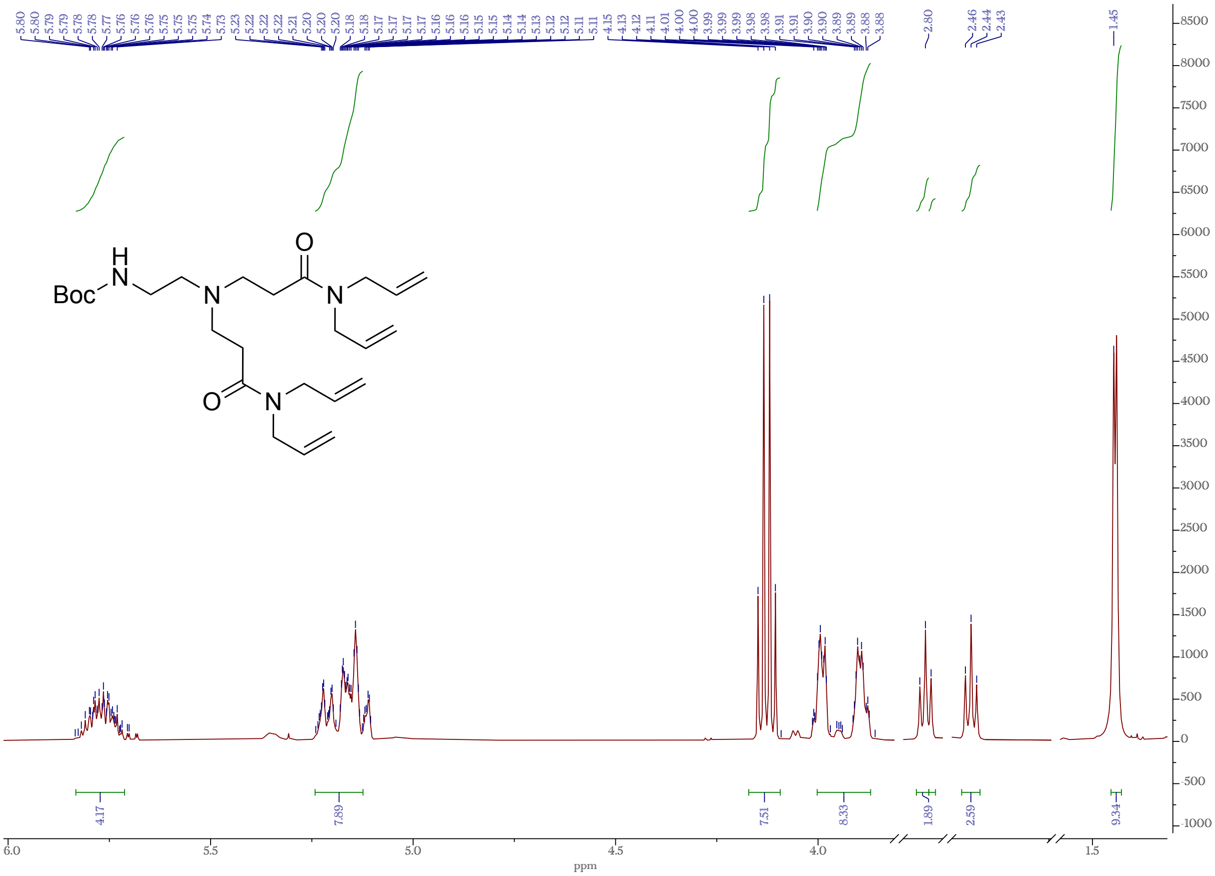

**Supplementary Figure S14.** $^{1}H$ NMR (500 MHz, $CDCl_3$) of *N*-Boc-bis(*N,N*-diallylpropanamide)-ethylenediamine.

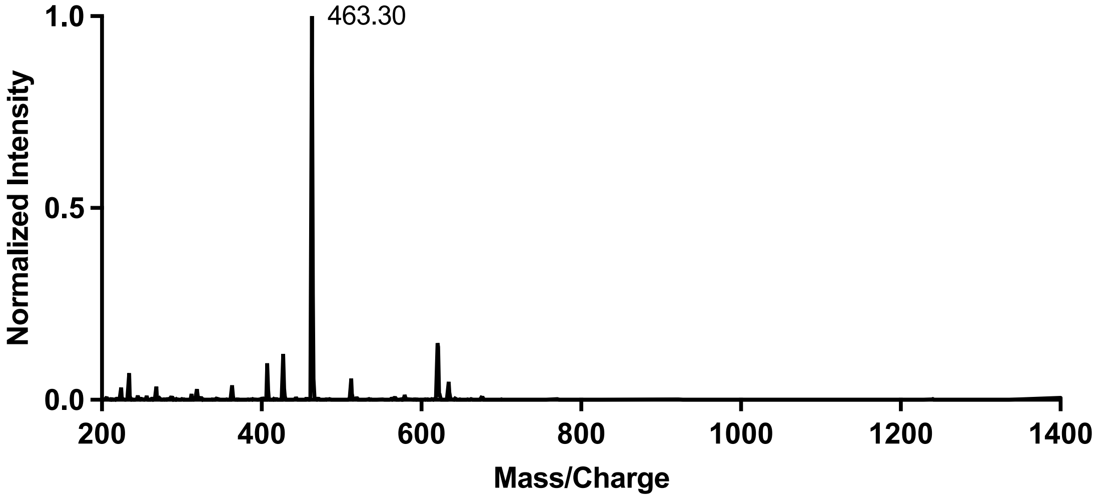

**Supplementary Figure S15.** LC/MS analysis of *N*-Boc-bis(*N,N*-diallylpropanamide)-ethylenediamine (positive ion mode).

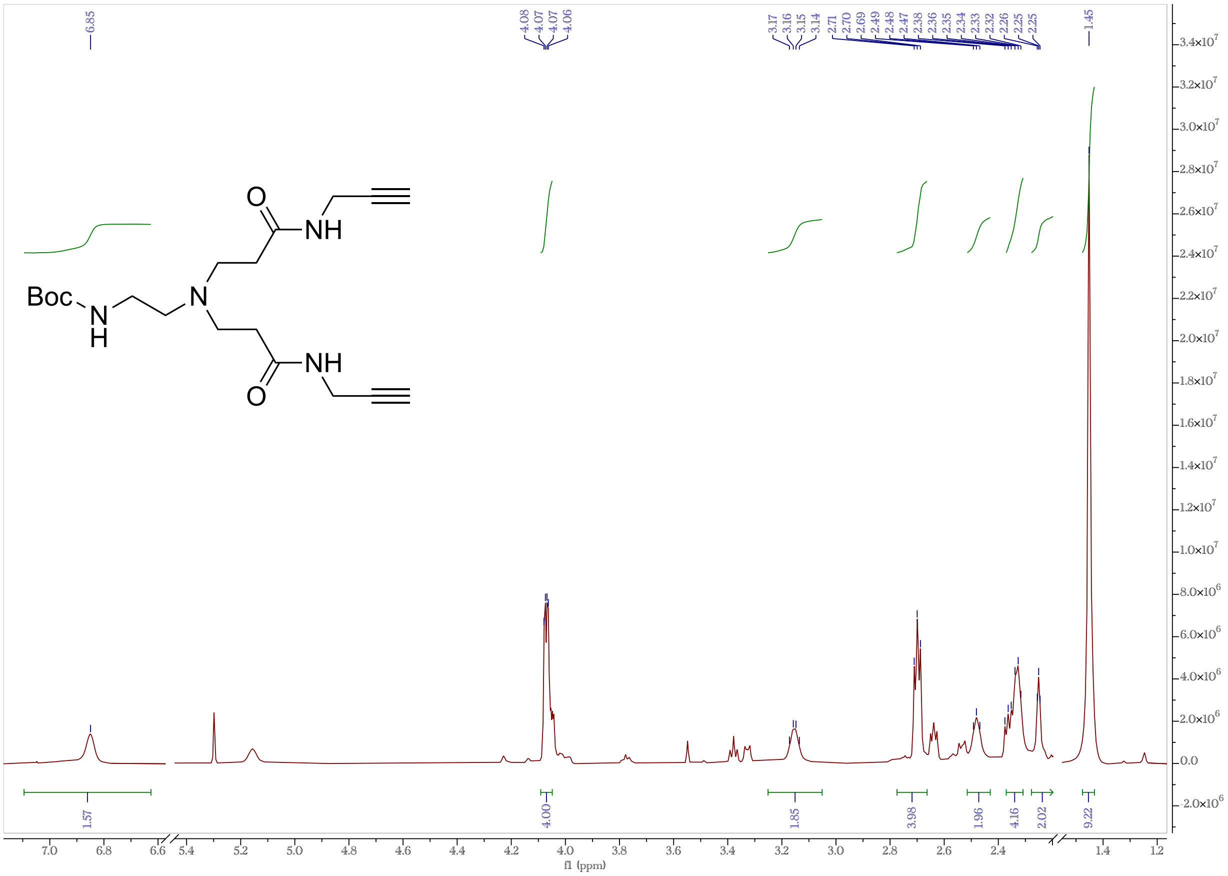

**Supplementary Figure S16.** $^1H$ NMR (500 MHz, $CDCl_3$) of *N*-Boc-bis(*N*-(prop-2-ynyl)lpropanamide)-ethylenediamine.

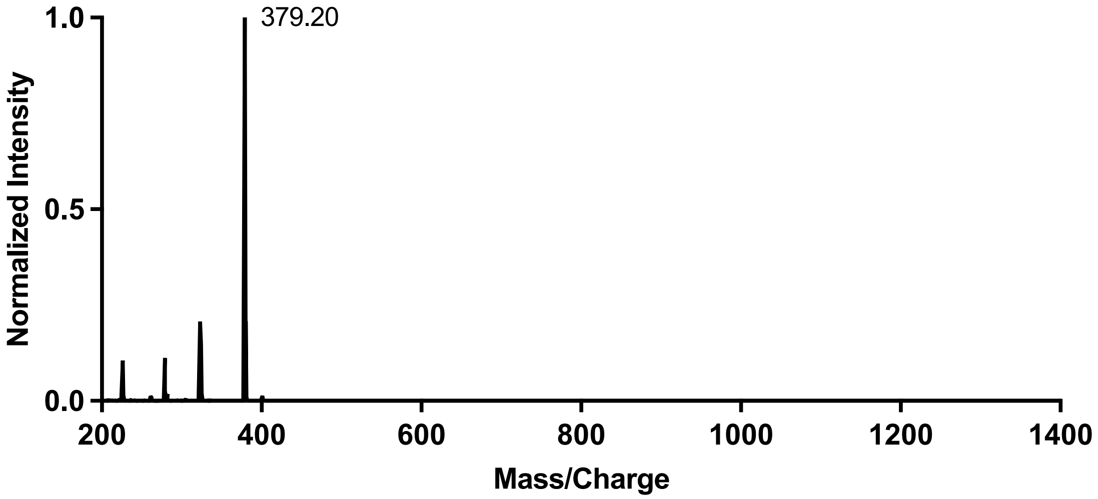

**Supplementary Figure S17.** LC/MS analysis of *N*-Boc-bis(*N*-(prop-2-ynyl)lpropanamide)-ethylenediamine (positive ion mode).

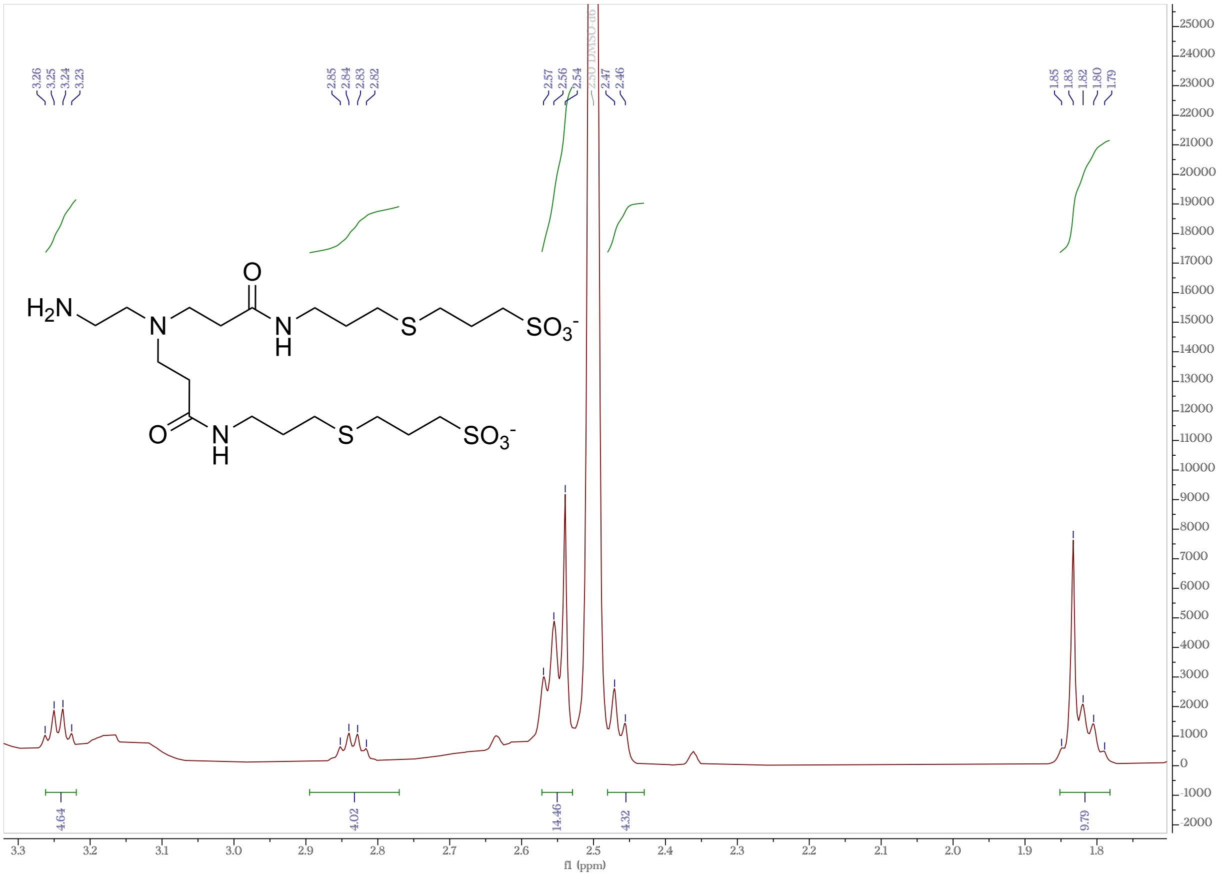

**Supplementary Figure S18.** $^1H$ NMR (500 MHz, DMSO-$d_6$) of amine-terminated SL2a.

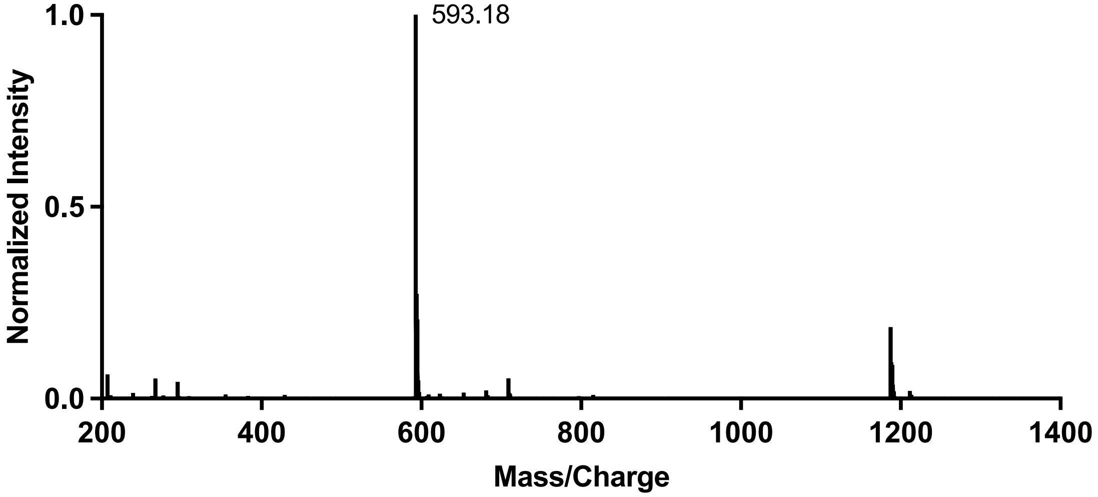

**Supplementary Figure S19.** LC/MS analysis of amine-terminated SL2a (negative ion mode).

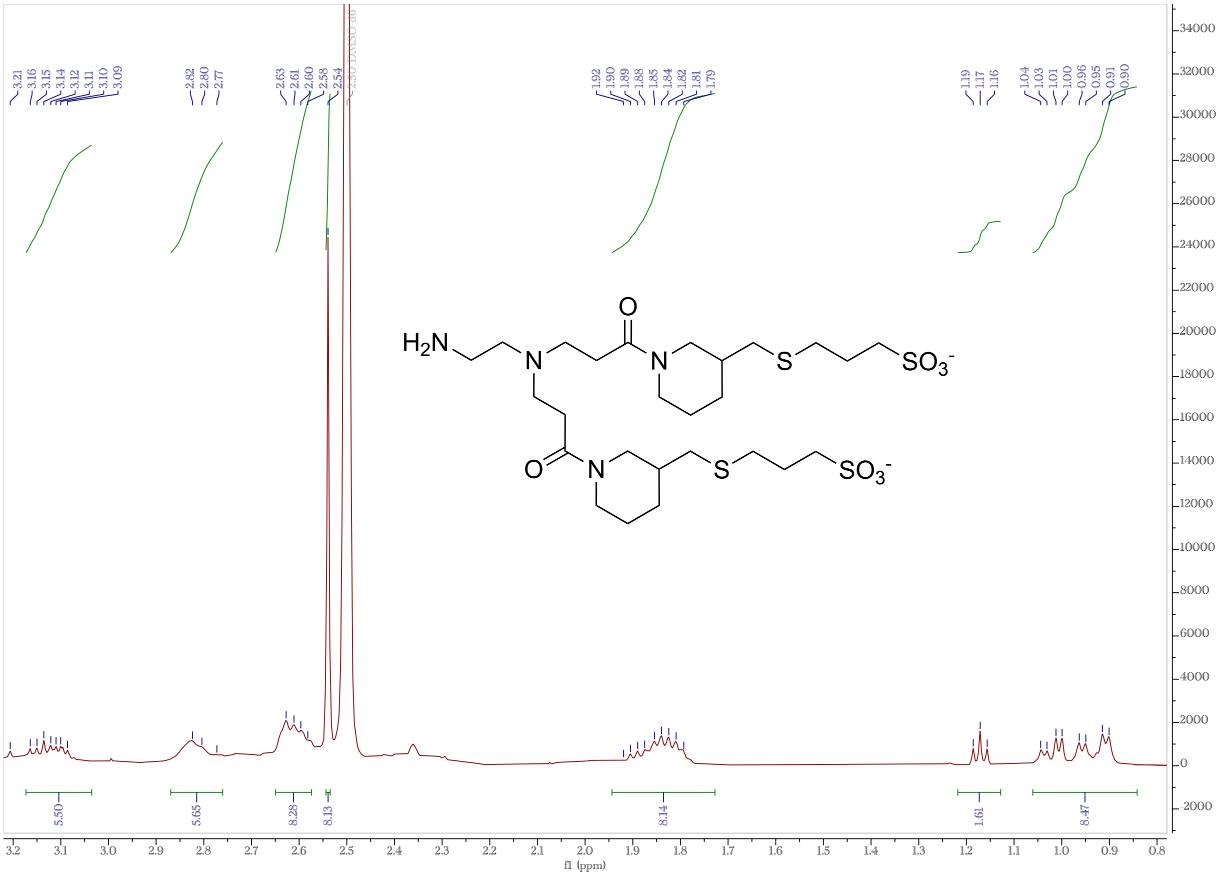

**Supplementary Figure S20.** $^1H$ NMR (500 MHz, DMSO-$d_6$) of amine-terminated SL2b.

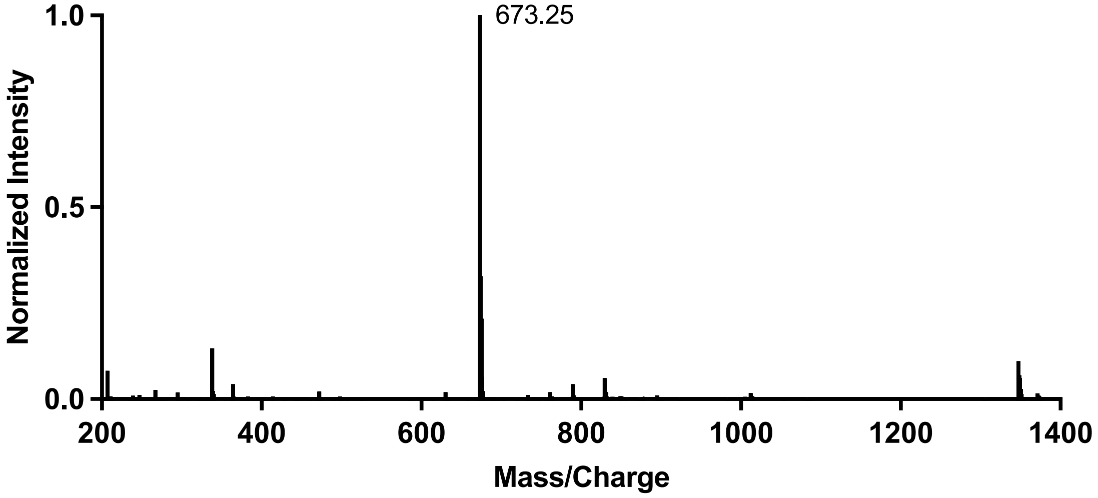

**Supplementary Figure S21.** LC/MS analysis of amine-terminated SL2b (negative ion mode).

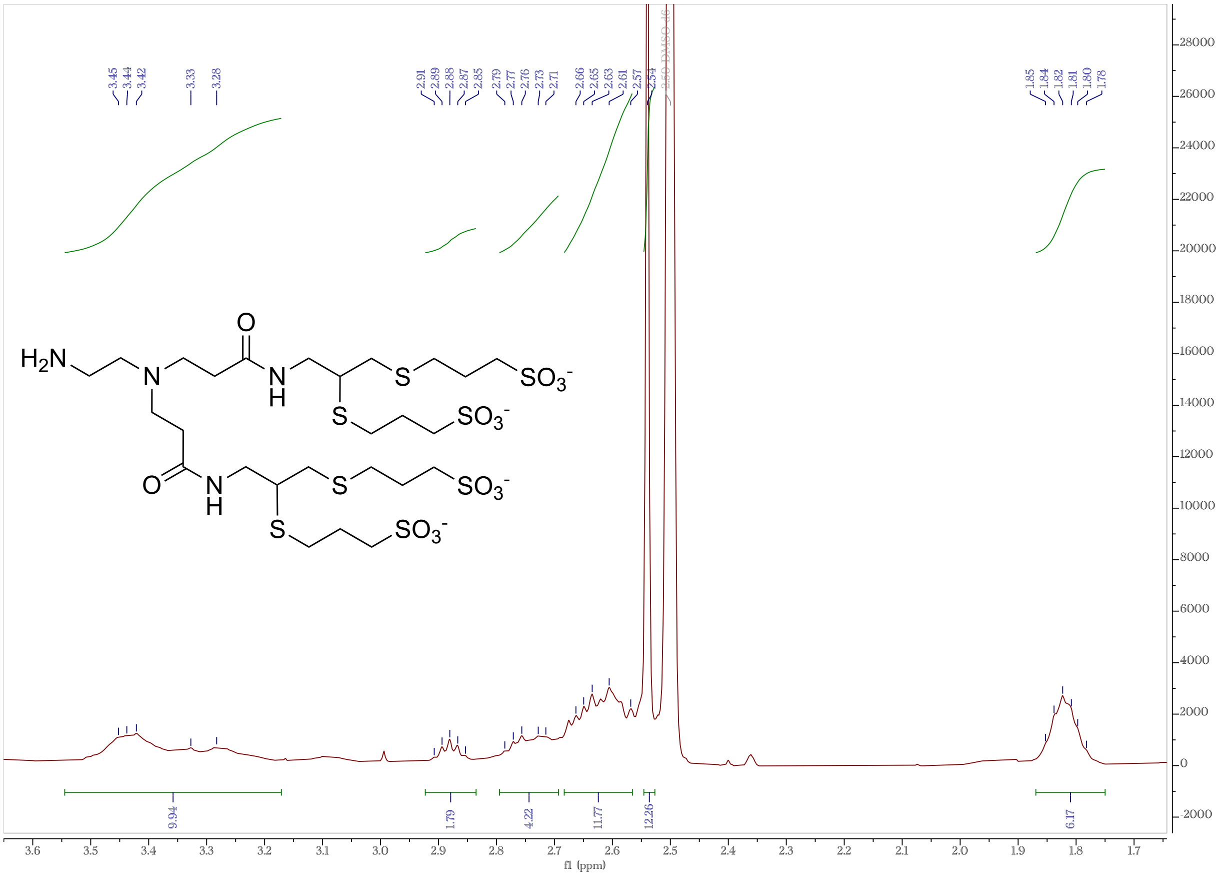

**Supplementary Figure S22.** $^{1}H$ NMR (500 MHz, DMSO-$d_6$) of amine-terminated SL4.

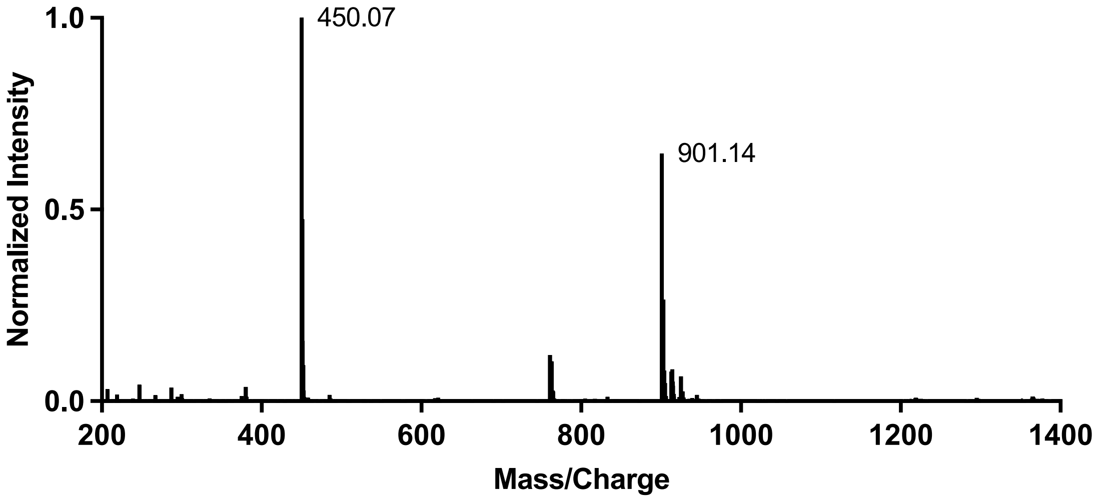

**Supplementary Figure S23.** LC/MS analysis of amine-terminated SL4 (negative ion mode).

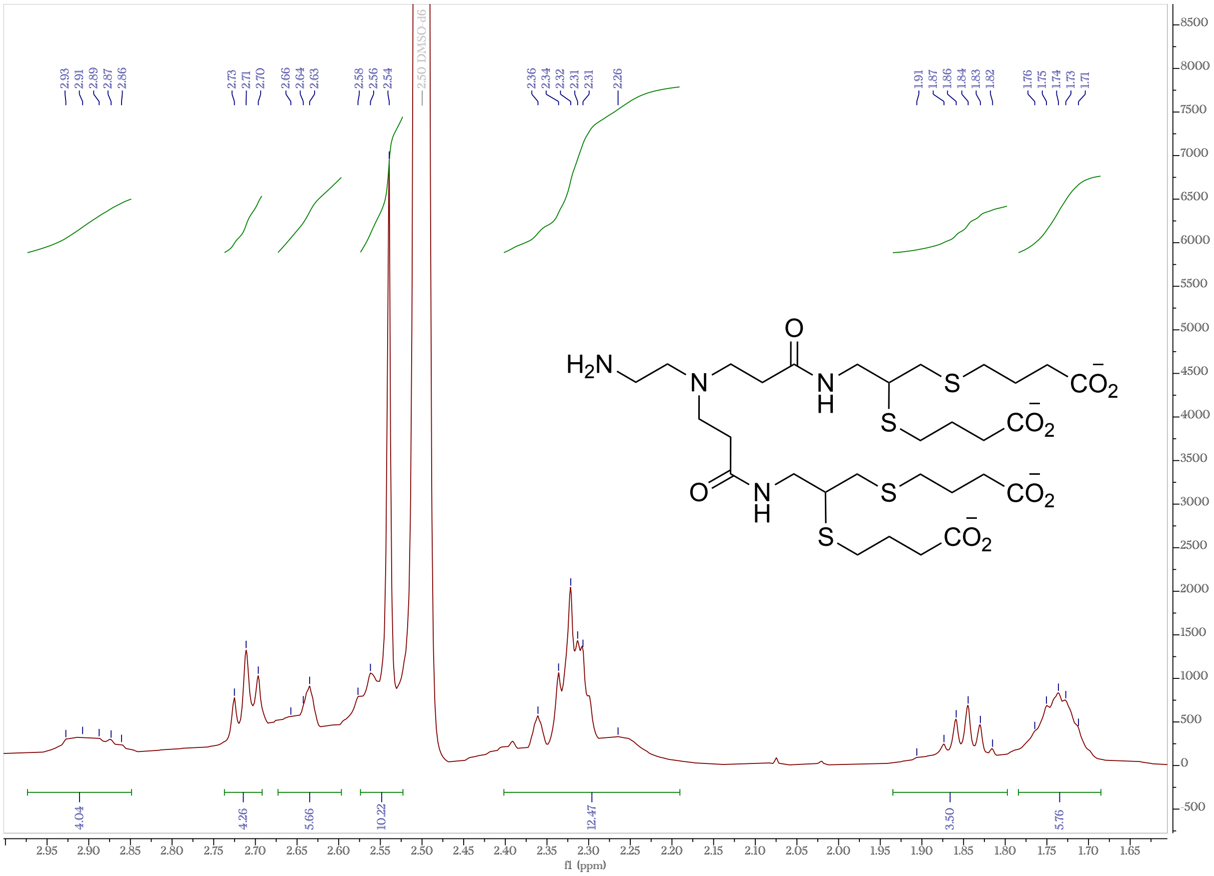

**Supplementary Figure S24.** $^{1}H$ NMR (500 MHz, DMSO-$d_6$) of amine-terminated CL4.

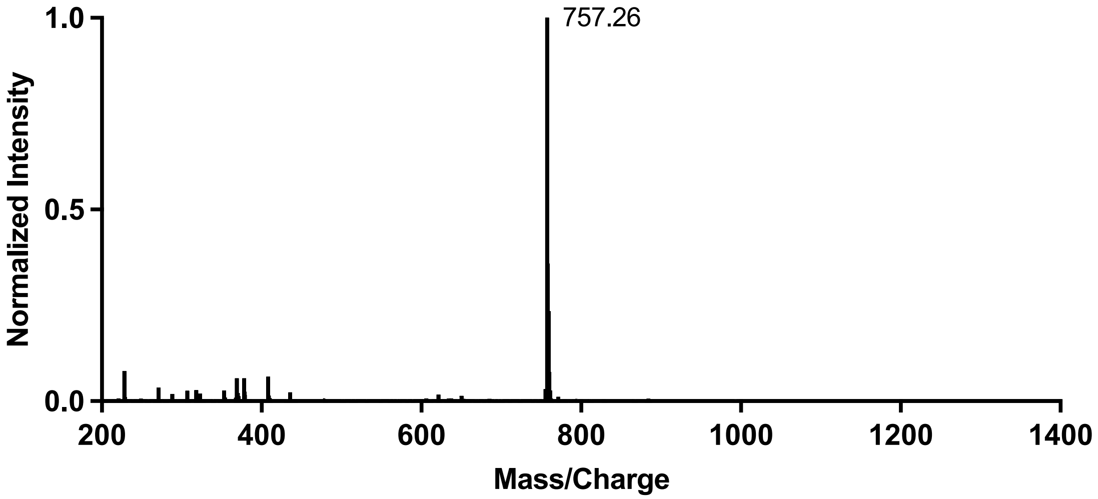

**Supplementary Figure S25.** LC/MS analysis of amine-terminated CL4 (negative ion mode).

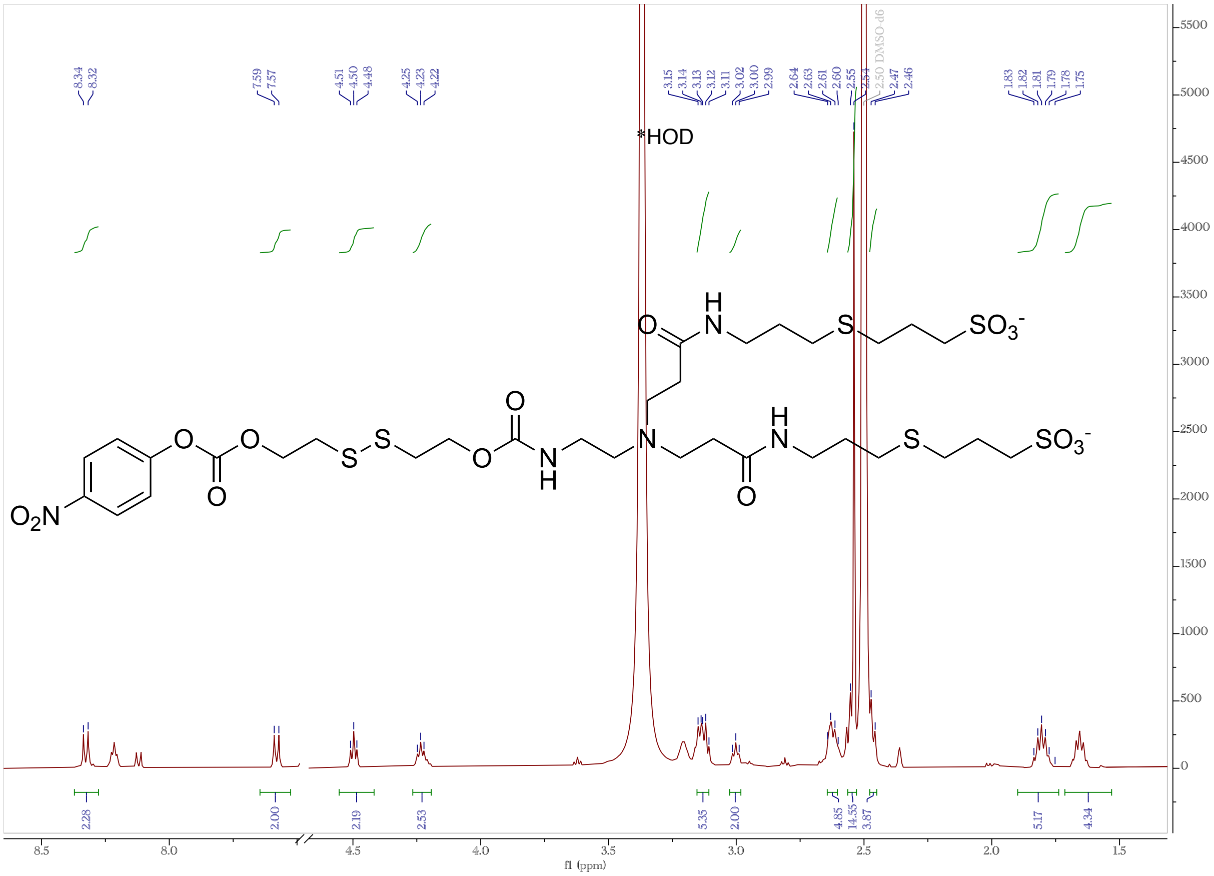

**Supplementary Figure S26.** $^1H$ NMR (500 MHz, DMSO-$d_6$) of SL2a.

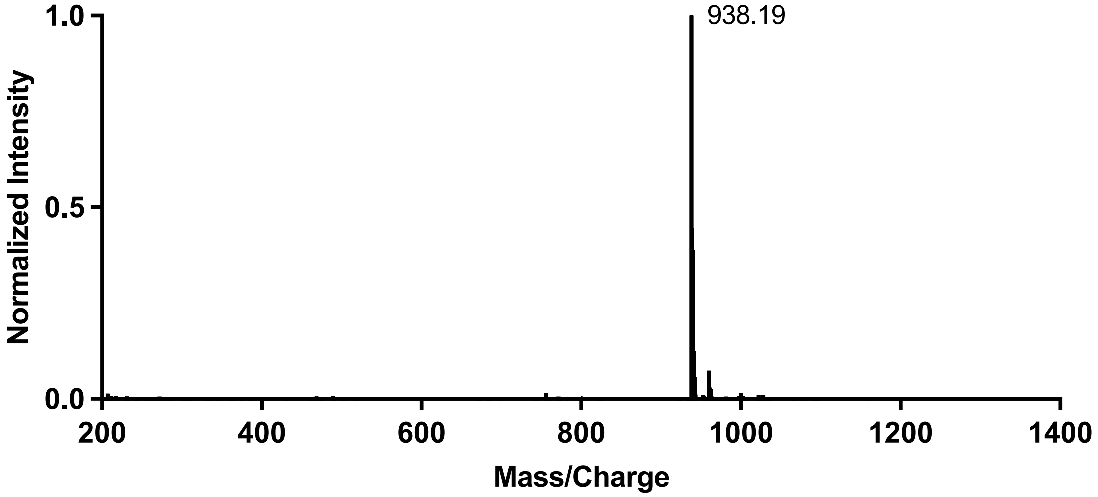

**Supplementary Figure S27.** LC/MS analysis of SL2a (negative ion mode).

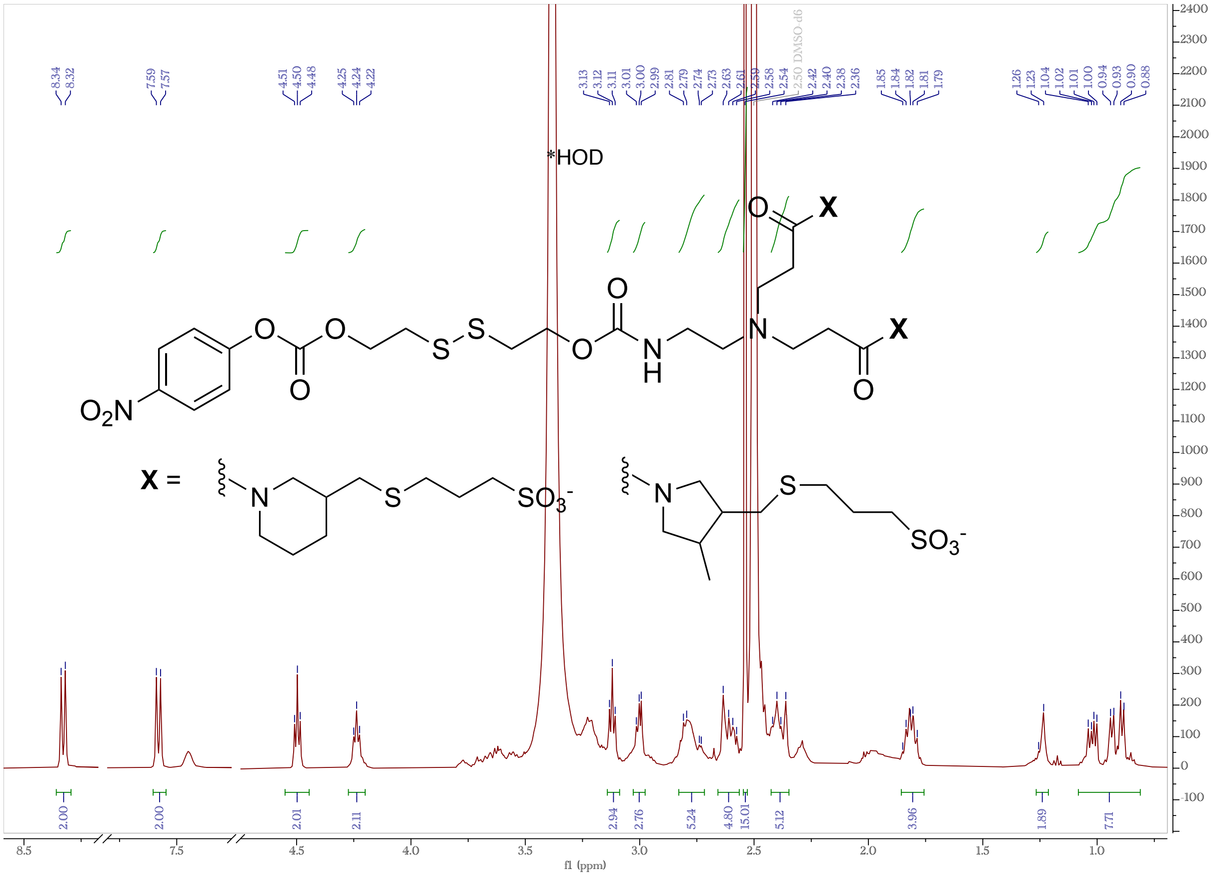

**Supplementary Figure S28.** $^1H$ NMR (500 MHz, DMSO-$d_6$) of SL2b.

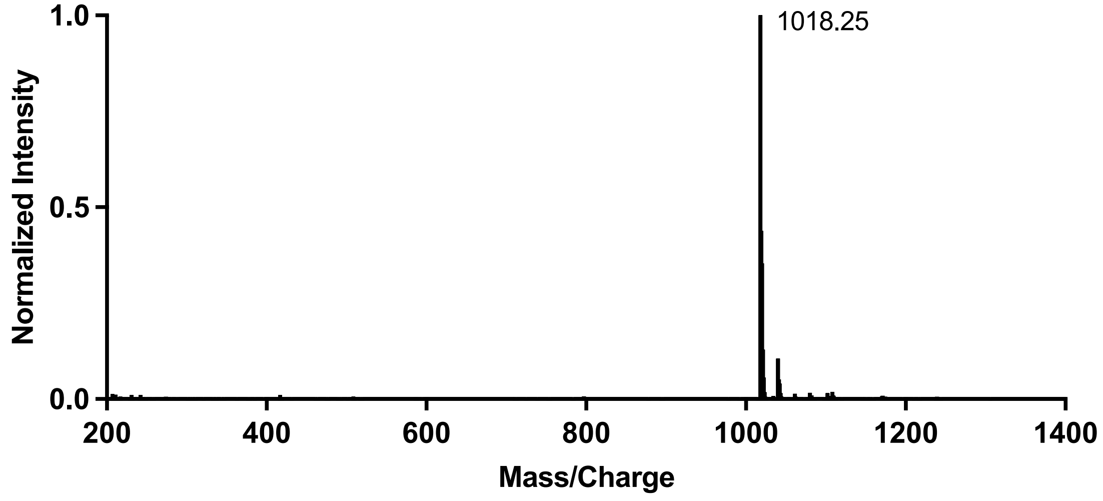

**Supplementary Figure S29.** LC/MS analysis of SL2b (negative ion mode).

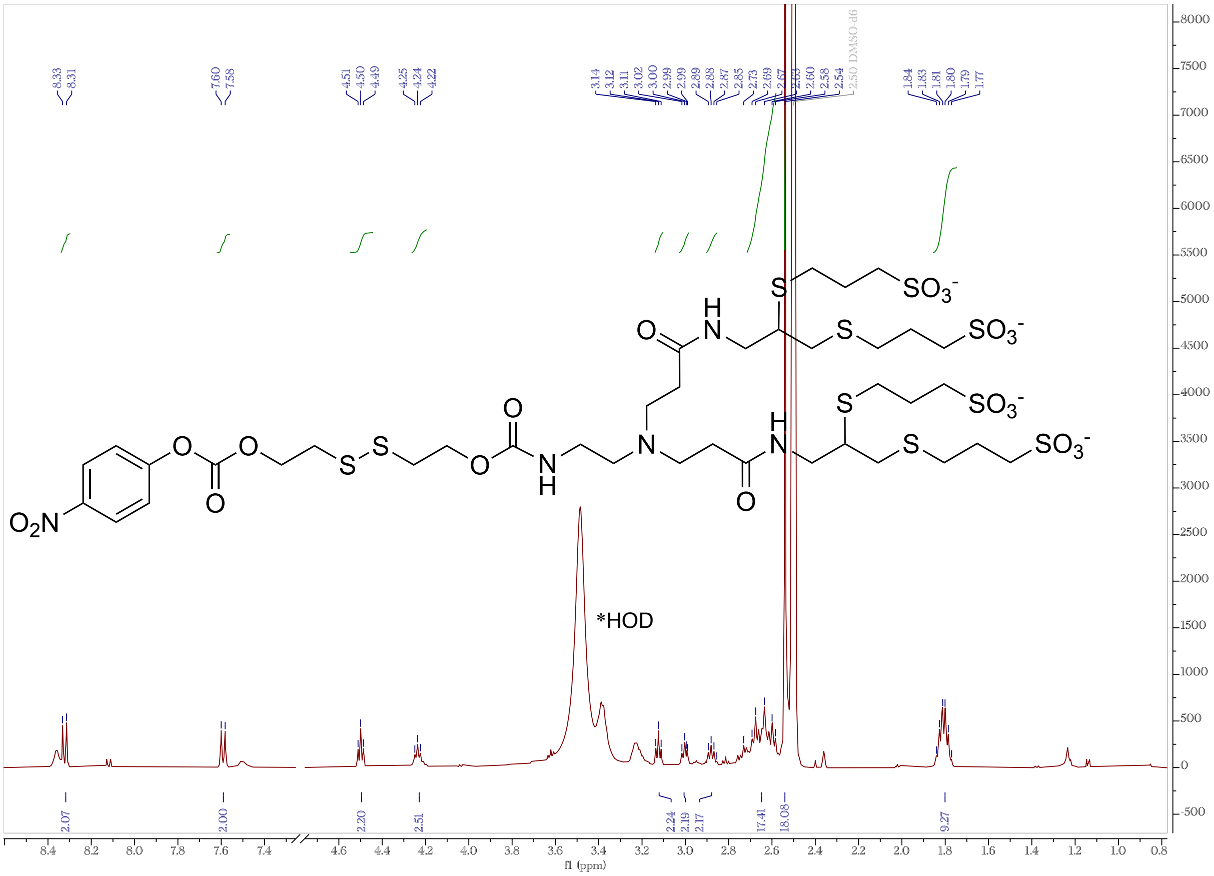

**Supplementary Figure S30.** $^1H$ NMR (500 MHz, DMSO-$d_6$) of SL4.

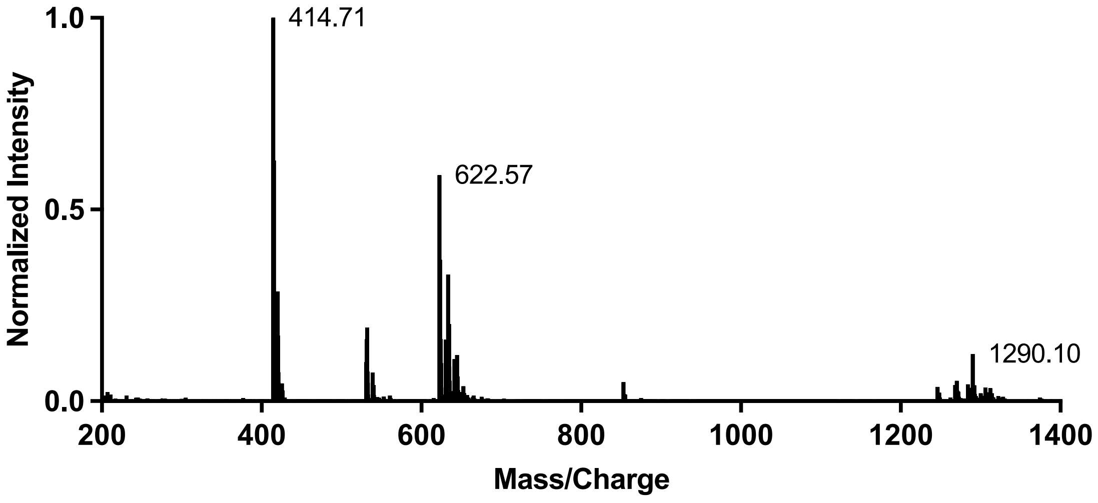

**Supplementary Figure S31.** LC/MS analysis of SL4 (negative ion mode).

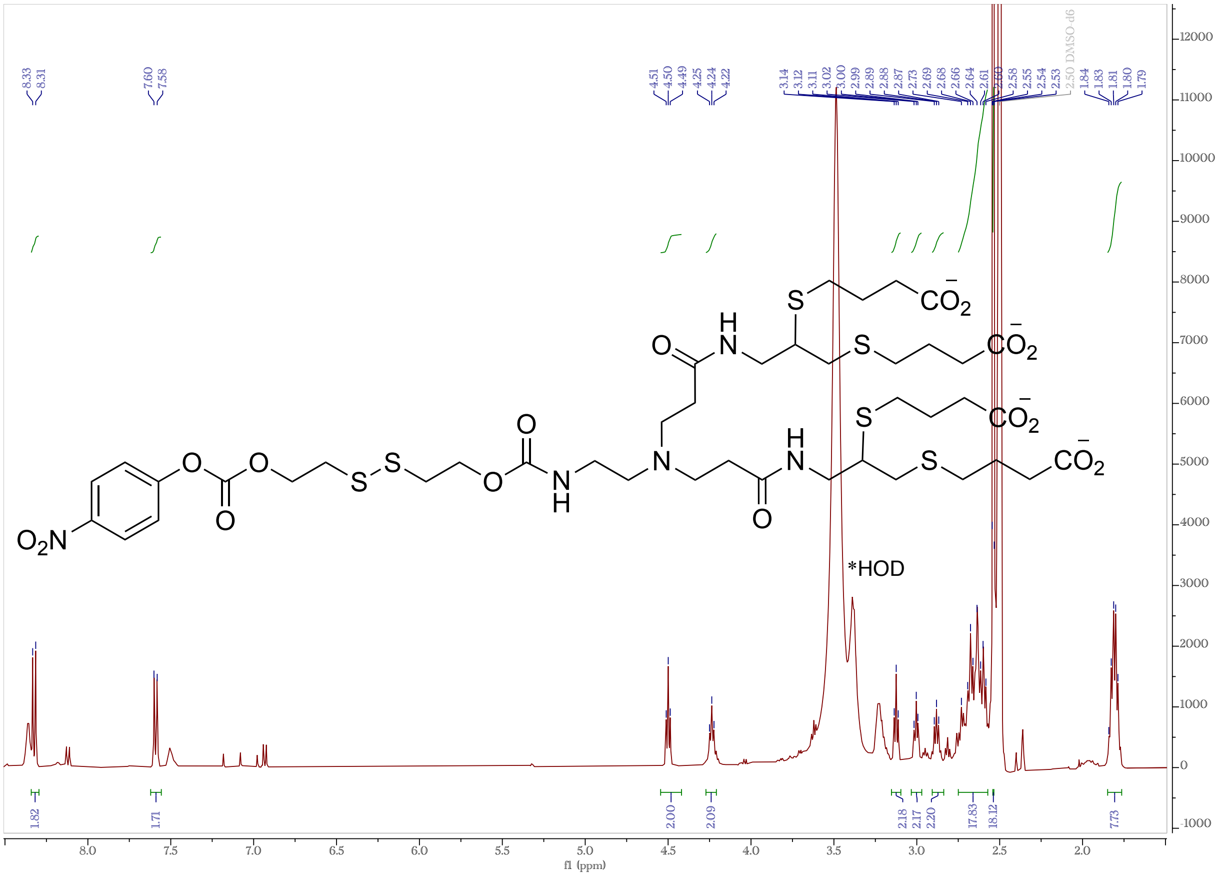

**Supplementary Figure S32.** $^1H$ NMR (500 MHz, DMSO-$d_6$) of CL4.

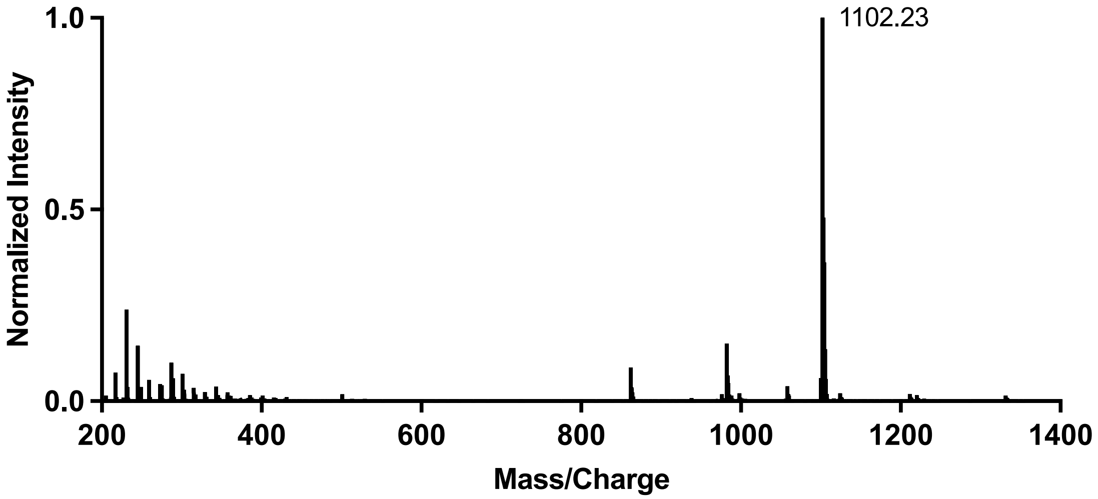

**Supplementary Figure S33.** LC/MS analysis of CL4 (negative ion mode).
